# Anatomic development of the upper airway during the first five years of life: A three-dimensional imaging study

**DOI:** 10.1101/2021.10.26.465942

**Authors:** Ying Ji Chuang, Seong Jae Hwang, Kevin A. Buhr, Courtney A. Miller, Gregory D. Avey, Brad H. Story, Houri K. Vorperian

## Abstract

**Purpose:** Normative data on the growth and development of the upper airway across the sexes is needed for the diagnosis and treatment of congenital and acquired respiratory anomalies and to gain insight on developmental changes in speech acoustics and disorders with craniofacial anomalies.

**Methods:** The growth of the upper airway in children ages birth-to-five years, as compared to adults, was quantified using an imaging database with computed tomography studies from typically developing individuals. Methodological criteria for scan inclusion and airway measurements included: head position, histogram-based airway segmentation, anatomic landmark placement, and development of a semi-automatic centerline for data extraction. A comprehensive set of 2D and 3D supra- and sub-glottal measurements from the choanae to tracheal opening were obtained including: naso-oro-laryngo-pharynx subregion volume and length, each subregion’s superior and inferior cross-sectional-area, and antero-posterior and transverse/width distances.

**Results:** Growth of the upper airway during the first five years of life was more pronounced in the vertical and transverse/lateral dimensions than in the antero-posterior dimension. By age five years, females have larger pharyngeal measurement than males. Prepubertal sex-differences were identified in the subglottal region.

**Conclusions:** Our findings demonstrate the importance of studying the growth of the upper airway in 3D. As the lumen length increases, its shape changes, becoming increasingly elliptical during the first five years of life. This study also emphasizes the importance of methodological considerations for both image acquisition and data extraction, as well as the use of consistent anatomic structures in defining pharyngeal regions.

## Introduction

The upper airway, a virtual conduit as characterized by Marcus et al. [1], has an anatomic boundary defined by other tissues (bony, cartilaginous and soft) while serving the functions of respiration, food ingestion (mastication and deglutition), as well as vocalization/speech, hence the function-based terms ‘*aerodigestive tract*’, ‘*vocal tract*’, or more comprehensively the ‘*aerodigestive and vocal tract’.* During the course of development, especially from infancy to early childhood, the upper airway undergoes drastic changes in size, shape and mechanical properties due to the restructuring of its anatomical sub-components, such as the descent of the larynx and the hyoid bone [2–4]. The anatomic growth process persists while adapting to the various functional needs and demands during maturation. As posited by current theory on craniofacial growth, the development of the upper airway is shaped by both genetic as well as intrinsic and extrinsic epigenetic factors, such as function, mechanical forces, and trauma [5–13].

The lack of knowledge regarding the growth and development of the upper airway, defined as the air conduit from the level of the nose to the carina, was addressed in a workshop by the National Heart, Lung, and Blood Institute (NHLBI) in 2009 with a large team of clinicians and scientists from diverse fields in healthcare and the biological sciences [1]. The outcome was a comprehensive set of research guidelines on various aspects of the upper airway, each with a set of priorities relevant to clinical disorders of upper airway functions. Among the priorities was the need to study the developmental changes of the upper airway anatomy and function during childhood (neonatal to puberty) across sexes and ethnicities and to provide normative values of the upper airway. Normative data are needed to better understand common respiratory disorders such as obstructive sleep apnea syndrome (OSAS), as well as a number of other congenital and acquired respiratory anomalies [1]. Furthermore, normative data can provide additional insight on developmental speech acoustics [14, 15], as well as speech disorders, particularly where craniofacial anomalies are present [16, 17]. As listed in Table 1, a large number of studies have examined the upper airway anatomy using different modalities, methodologies, airway regions, and age ranges. Table 1 summarizes the studies to date that have examined the typical development of the aerodigestive and vocal tract from the choanae or the soft palate superiorly to the epiglottis or the trachea inferiorly. A subset of studies listed have factored in growth and/or sex in their data analysis. Most studies have employed imaging to obtain quantitative measurements, including linear, angular and/or area measurements, based on the midsagittal or axial slices, as well as volumetric measurements. However, only a very limited number of studies have assessed multidimensional volumetric measurements during early childhood. Among the 34 studies listed in Table 1, only 17 studies included linear, area and volumetric measurements, and fewer than half of those studies controlled for head position during or after data acquisition. Of the 12 studies summarized in Table 1 that examined the pre-pubertal age-group, the majority obtained measurements in 2D that were collected primarily from radiographic images using mid-sagittal, axial, or coronal visualization planes; an approach frequently used to assess the upper airway, as it is cost effective and less time-consuming to process. However, this approach does not provide accurate representation of the complex airway morphology, as it overlooks information of lateral dimensions [18, 19]. Two of those 12 studies [20, 21] quantified the prepubertal airway in 3D but only Abramson et al. [20] covered the entire prepubertal period from birth to five years and assessed sexual dimorphism. Neither of those retrospective studies reported controlling for head position or using it as an inclusion criterion.

**Table 1.**
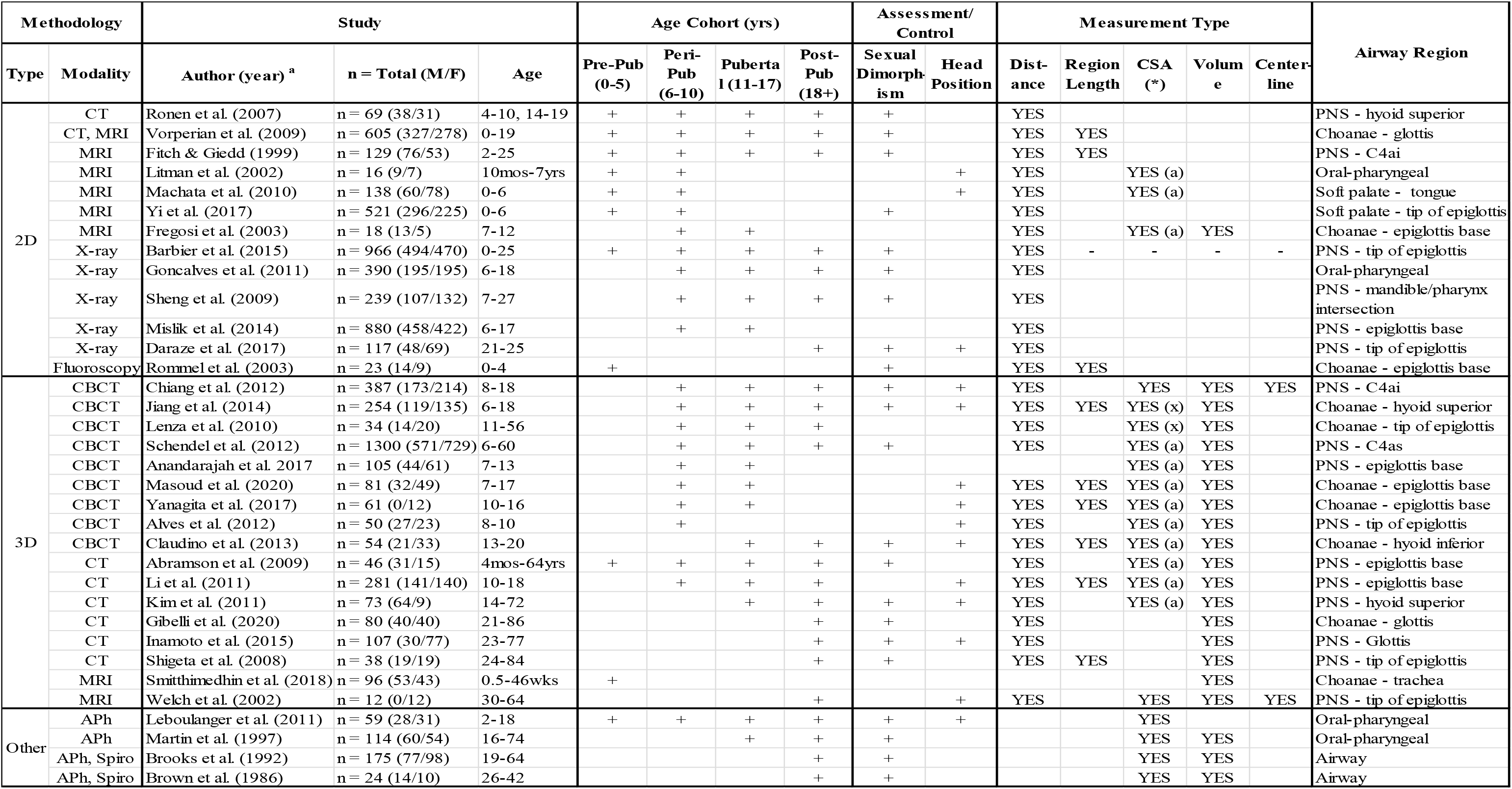
Summary of studies on typical upper airway development. Summary of studies on typical upper airway development listed per methodology in the **first** column, using measurement type/dimension: 2D (two-dimensional), indicates that measurements were collected from radiographic images representing either the mid-sagittal, axial or coronal visualization planes; 3D (three-dimensional), indicates that measurements were collected utilizing multiplanar visualizations (axial, coronal and/or sagittal planes) and 3D representation of the upper airway; Other indicates that measurements were made using non-imaging techniques. The first column also lists alphabetically study modality including imaging (CT, CBCT, MRI, X-ray) or non-imaging (Acoustic Pharyngometry (APh), Spirometer (Spiro)) techniques. The **second** column lists alphabetically the study author(s) with year of publication in parentheses, sample size (n=) with total male/female (M/F) numbers specified in parentheses, and age range examined. Age of study participants is also classified using pubertal age cohorts in the **third** column, followed by assessment or control of sex-differences and head position in the **fourth** column. The **fifth** column lists measurement reported including overall pharyngeal distance, and pharyngeal region/subregion measurements including: length, cross-sectional area* (CSA), volume and centerline length. The **final** column lists the defined superior to inferior anatomical boundaries in each study. * CSA specifications include the automatic or manual calculation of the area of a plane of the pharyngeal airway in: the axial plane (a), a plane that is orthogonal to the pharyngeal centerline (o), or a plane connecting two or more specific anatomical landmarks (x). ^a^ Cited literature: [20–53]

Since the upper airway is a lumen, attention must be paid to a number of methodological considerations, given their potential effect on various pharyngeal measurements. Methodological procedures known to affect pharyngeal measurements include head/neck position (flexion/extension), body position (upright/supine), and sedation [37, 54, 55]. While most studies to date have accounted for one or more of these confounders, it is difficult to compare findings across studies unless all confounders have been addressed. For example, Inamoto et al. [47], reported significant sex differences between the adult male and female laryngopharynx, but Gibelli et al. [46], who also used CT but did not control for head position, reported no sex differences. Additionally, variations in the anatomical boundary and subregion borders of the upper airway morphology (nasopharynx, oropharynx, and laryngopharynx/hypopharynx) as defined by different studies, summarized in Table 1 (final column) and Table 2, further complicates the ease and feasibility of comparing findings across studies. Thus, to ensure an accurate and reliable assessment of the developmental changes of this cavity, it is critical to use standardized imaging procedures, well-defined anatomical regions, and established airway data extraction protocols.

**Table 2.** Summary of boundaries of the pharyngeal regions. Summary of pharyngeal regions’ boundaries as defined in relevant anatomy textbooks and published papers. Note the lack of consistency and/or specificity in the anatomical boundaries for each region. ^b^ Cited literature: [56–65]

| Main Pharyngeal Subdivisions | Anatomical boundaries | Definitions <sup>b</sup> |
| --- | --- | --- |
| Nasopharynx | **Superior Border** | Nares (Ayappa & Rapoport, 2002)<br>Nasal cavity (Adewale, 2009)<br>End of nasal septum (Netter, 2019)<br>Choanae (Arens et al., 2004; Logan et al., 2017; Schuenke et al., 2010) |
|  | **Inferior Border** | Hard palate (Ayappa & Rapoport, 2002)<br>Soft palate (Laird et al., 2019 ; Moore et al., 2006; Standring et al., 2017)<br>Level of soft palate (Arens et al., 2004)<br>Above soft palate (Adewale, 2009; Gu et al., 2016 )<br>Inferior/Lower border of soft palate (Netter, 2019; Logan et al., 2017 ) |
| Oropharynx | **Superior Border** | Soft palate (Ayappa & Rapoport, 2002)<br>Uvula (Schuenke et al., 2010) |
|  | **Inferior Border** | Epiglottis (Ayappa & Rapoport, 2002; Schuenke et al., 2010)<br>Superior/Upper border of epiglottis (Gu et al., 2016; Laird et al., 2019 ; Moore et al., 2006; Netter, 2019; Standring et al., 2017)<br>Larynx (Arens et al., 2004) |
| Laryngopharynx/<br>Hypopharynx | **Superior Border** | Posteriorlateral to the larynx (Arens et al., 2004)<br>Base of tongue (Ayappa & Rapoport, 2002)<br>Tip of epiglottis (Adewale, 2009)<br>Superior/Upper border of epiglottis (Logan et al., 2017 ) |
|  | **Inferior Border** | Larynx (Ayappa & Rapoport, 2002)<br>Opening of esophagus (Netter, 2019)<br>Cricoid cartilage (Schuenke et al., 2010)<br>Inferior/Lower border of cricoid cartilage (Adewale, 2009; Gu et al., 2016; Laird et al., 2019; Logan, 2017; Moore et al., 2006; Standring et al., 2017) |

This study aims to systematically study the anatomic development of the upper airway, specifically the structural changes from the choanae to the tracheal opening (inferior border of the cricoid cartilage) from birth to five years, as compared to adults. To acquire normative data of the anatomy that subserve the aerodigestive and speech functions, we used an imaging database with computed tomography (CT) studies from typically developing individuals to obtain a comprehensive set of two-dimensional (2D) and three-dimensional (3D) measurements quantifying the growth of the upper airway. Our comprehensive set of methodological criteria included control of head position, histogram-based upper airway segmentation, placement of anatomic landmarks, and development of a semi-automatic method to determine lumen centerline. In addition, to better understand the resonance/acoustic characteristics of the vocal tract, this study aimed to examine the nature of the developmental changes of the upper airway dimensions and to determine if there are sex differences in the upper airway dimensions during the pre-pubertal period. We hypothesize all pediatric airway dimensions to be substantially smaller than adult dimensions. We also hypothesize sex differences in both children and adults.

## Materials and methods

### a. Image acquisition/dataset

Using imaging studies performed at the University of Wisconsin Hospital and Clinics (UWHC), our Vocal Tract Development Lab (VTLab) has curated a lifespan retrospective database of more than 2000 head and neck CT scans to study the anatomic growth and development of the oral and pharyngeal structures. This database was established following approval of the University of Wisconsin-Madison Institutional Review Board (IRB) and anonymized accordingly. All CT imaging studies, performed in the supine body position, were acquired using CT scanners manufactured by General Electric Medical Systems or Siemens and stored in Digital Imaging and Communications in Medicine (DICOM) format. Additional details on this imaging database and image acquisition are provided in Kelly et al. [66], Miller et al. [67, 68] and Vorperian et al. [23].

To ensure the adequacy of imaging studies selected for this study, the VTLab imaging database was reviewed for typically developing cases between the ages 0-5 years (pediatric) and 20-30 years (adults) who were imaged for conditions that do not affect typical growth. A total of 410 (208 Males (M), and 202 Females (F)) CT imaging studies that included 264 (161M, 103F) pediatric scans and 146 (47M, 99F) adult scans, from 276 (143M, 133F) individuals (195 [115M, 80F] children; and 81 [28M, 53F] adults), were inspected for cases that met the following inclusion criteria: (1) slice thickness ≤ 2.5mm, (2) 14-22cm field-of-view (FOV), (3) 512x512 matrix size, (4) no movements or dental artifacts affecting the view of pharynx structure, and (5) neutral or flexed head position as confirmed using Miller et al.’s [68] head position classification protocol. While all extreme flexion/extension cases were excluded, including all sedation cases, neutral-flexed head position cases were not excluded given that the larger infant head is prone to being flexed in the supine position. The total yield of cases that met the inclusion criteria for this study’s dataset included 61 (32M, 29F) pediatric cases from 78 imaging studies (41M, 37F), and 17 (9M, 8F) adult cases from 72 (39M, 33F) imaging studies. The individuals whose images were used included 56 (31M, 25F) children, and 16 (8M, 8F) adults. Age specific demographics are presented in Table 3.

**Table 3.** Distribution of male and female cases per age group. Distribution of male (M) and female (F) cases per age group. Age groups specified in years; months (group <1 includes cases birth (00;00) to 11 months (00;11); group 1 includes cases 1 year (01;00) to 1 year 11 months (01;11) etc., and group 5 adults ages 20-to-30 years.

| Group (Age range (yr;mos)) | M | F | Total |
| --- | --- | --- | --- |
| <1 (00;00 - 00;11) | 4 | 5 | 9 |
| 1 (01;00 - 01;11) | 4 | 5 | 9 |
| 2 (02;00 - 02;11) | 6 | 7 | 13 |
| 3 (03;00 - 03;11) | 9 | 8 | 17 |
| 4 (04;00 - 04;11) | 6 | 7 | 13 |
| 5 (20;00 - 30;00) | 8 | 9 | 17 |

### b. Image reconstruction

The standard reconstruction kernel was the preferred CT reconstruction algorithm, and was available for the majority of the imaging studies. For cases/imaging studies processed without the standard kernel, imaging features of the standard kernel were simulated by processing the soft kernel with an unsharp enhance filter using a kernel size of 5x5, or the by processing the bone kernel with a low pass filter using a kernel size of 3x3. Next, the software Analyze 12.0 [69] was used to reconstruct CT images from DICOM format into 3D volume.

A histogram-based threshold method was applied to the reconstructed CT volume in order to identify the intensity in Hounsfield Unit (HU) that allows an optimal representation of the airway. Guided by the technique of Nakano et al. [70], per image, we used the midpoint between the air threshold peak (-1000 HU) and soft tissue peak (+100 HU to +300 HU), as the applied upper threshold intensity to segment the airway. The range of upper thresholds used in this study was between -556 HU and -445 HU. The Volume Render and Volume Edit modules were then used to visualize and segment the 3D pharynx model from the reconstructed CT volume. Using the identified threshold value, the airway region studied was restricted inferiorly at the first tracheal ring (lower limit of the cricoid cartilage), and superiorly at the choanae. The resulting 3D pharynx model was saved in Analyze Object Map format [.obj].

### c. Anatomic landmarks and variables

As depicted in Figs 1 and 2, and listed with descriptions in Table 4, a set of 26 anatomic landmarks that included 20 pharyngeal, 4 maxillary, and 2 reference landmarks were manually placed on each of the 78 3D pharynx models to quantify upper airway growth. The set of landmarks selected were carefully determined following a thorough review of landmarks and airway variables examined in studies to date [20, 24, 36, 47, 54, 71]. Landmark placement entailed using the Volume Render module in Analyze 12.0 [69], to manually place each of the 26 landmarks by overlaying them on their respective CT images while using the axial, coronal and sagittal planes to guide accuracy of landmark placement. The landmarks were similarly saved in Analyze Object Map format. To ensure reliability in landmark placement, prior to data collection, two researchers modeled and placed landmarks on six cases. The differences in resulting measurements, calculated from the raters’ landmarks, had an average relative error (ARE) that was less than or equal to 5% between researchers. The landmarks were then used to establish a data extraction protocol, described in the following section, that generates pharyngeal cross sections perpendicular to the centerline and calculates landmark-based measurements. The comprehensive set of 30 pharyngeal variables measured, as listed and defined in Table 5 below, are described in the section on variable measurements.

**Fig 1.**
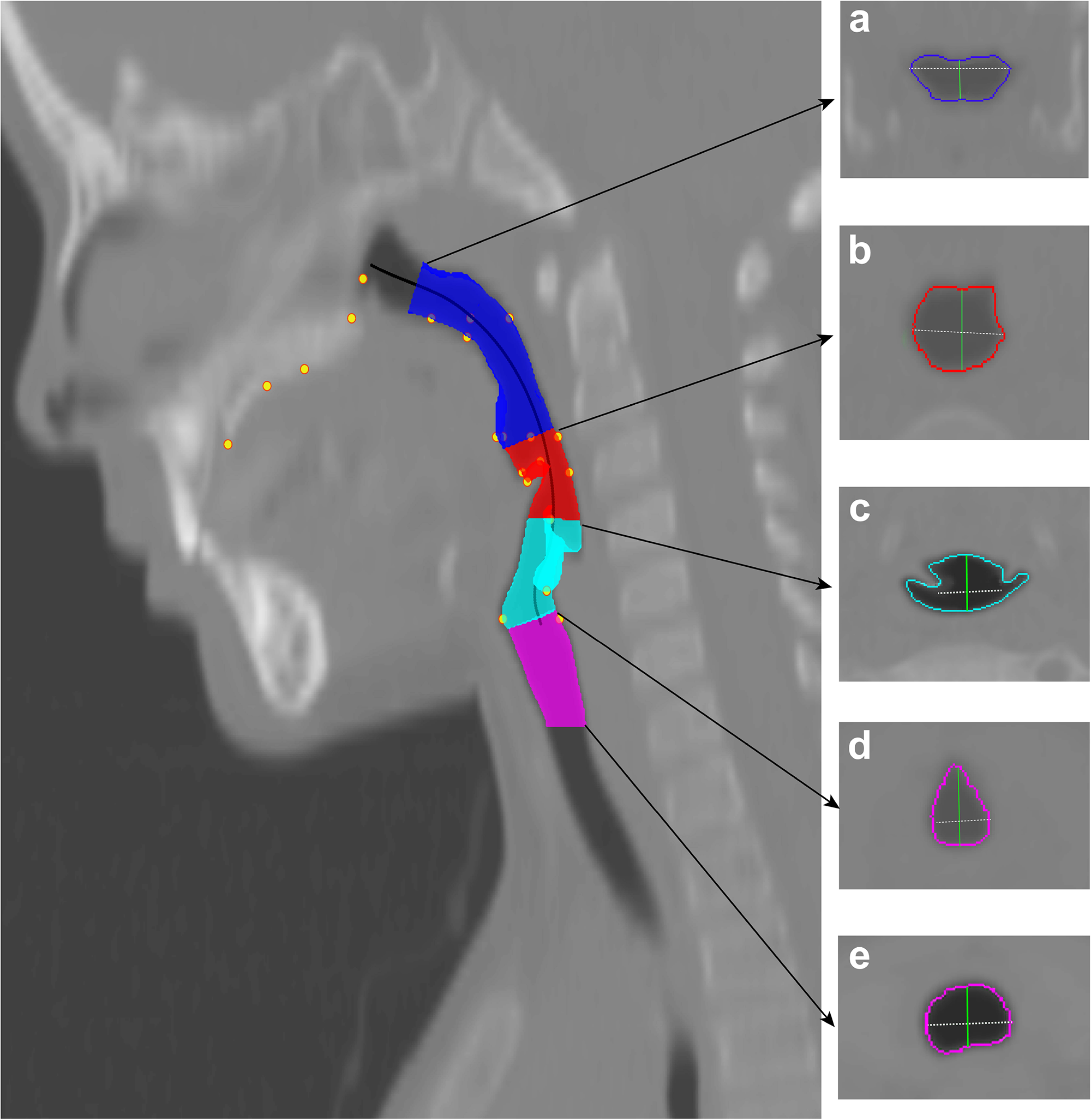
Illustration of airway regions and measurements. The airway was examined using landmark-derived planes orthogonal to the centerline, as described in text. The four airway regions bounded by five cross sectional areas, a-to-d as depicted in the right panel, were quantified developmentally using the following measurements: volume, region length, cross-sectional area (CSA), anterior-posterior distance, and lateral width - as defined in Table 5. The airway regions above the glottis (d; Table 5, definition 19), included the following pharyngeal regions: a. *Nasopharynx* (blue; definition 1); b. *Oropharynx* (red; definition 6); c. *Laryngopharynx* (cyan; definition 11); and the airway below the glottis consisted of the *subglottal* region (magenta; definition 22). The pharynx (Table 5, definition 16) consisted of all three supraglottal regions.

**Fig 2.**
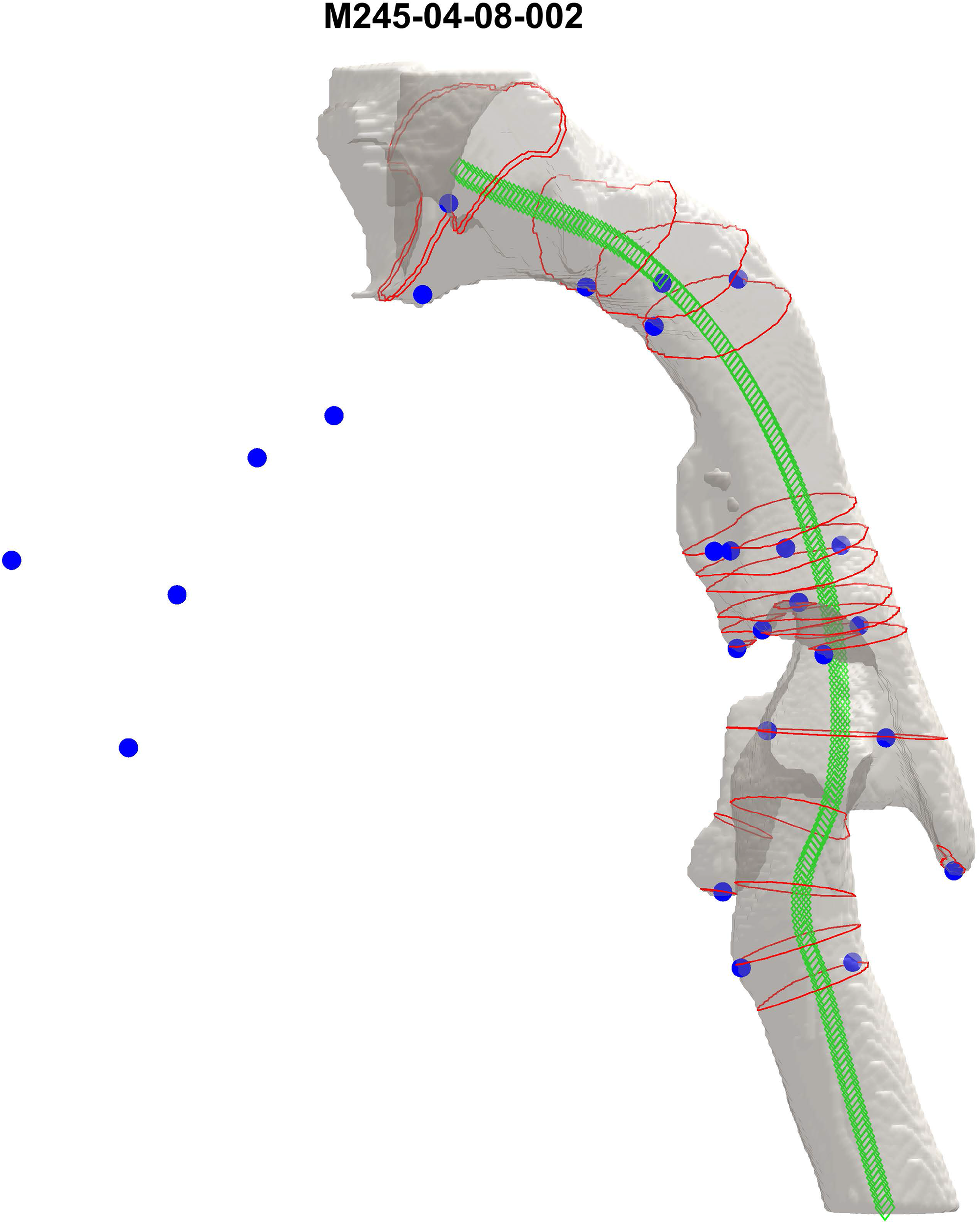
3D airway model (choanae to trachea) of a 4-year 8-month old typically developing male as visualized in MATLAB. Blue dots represent the 26 anatomic landmarks listed in Table 4. Centerline is shown in green, and CSAs closest to each of the anatomic landmarks are shown in red.

**Table 4.** Description/definition of the 26 anatomic landmarks. Description/definition of the 26 anatomic landmarks (pharynx, reference, and maxilla), listed from the inferior to the superior regions of the airway as displayed in Fig 1. Landmark placement entailed use of multiplanar views (at least two of the sagittal, axial, and coronal planes, or all three) for accuracy. These landmarks were used in defining study variables and extracting the quantitative measurements of the upper airway as specified in Table 5.

| # | Description of landmarks | Landmark Name | Abbreviation |
| --- | --- | --- | --- |
| **Pharynx Landmarks** |  |  |  |
| 1 | The point of attachment of the vocal folds with the thyroid cartilage at the anterior commissure, 2-3 mm below the thyroid notch of the larynx. | Glottis Anterior | (ga) |
| 2 | In the axial plane at the level of the glottis as determined by the tear-shaped glottal area (GA), the most posterior point of the laryngo-pharynx between the lateral-most sides of the vocal folds attached to the arytenoid cartilages. | Glottis Posterior | (gp) |
| 3 | Most superior and posterior point of the epiglottis in the midsagittal plane. | Epiglottis Superior | (epS) |
| 4 | Attachment of the epiglottis with the hyoepiglottic ligament, visualized as the most anteroinferior point of contact in the midsagittal plane. | Epiglottis Base | (epBse) |
| 5 | The most posterior point on the pharyngeal wall at the level of the epiglottis base in the midsagittal plane. | Epiglottis Base Posterior | (epBsePo) |
| 6 | Most inferior point of the left piriform sinus. | Piriform Sinus Inferior Left | (PSInL) |
| 7 | Most inferior point of the right piriform sinus. | Piriform Sinus Inferior Right | (PSInR) |
| 8 | The midpoint (most 'curved' point) of the left aryepiglottic fold . Approximately halfway between the base and tip of the epiglottis. | Piriform Sinus Superior Left | (PSSuL) |
| 9 | The midpoint (most 'curved' point) of the right aryepiglottic fold. Approximately halfway between the base and tip of the epiglottis. | Piriform Sinus Superior Right | (PSSuR) |
| 10 | Most inferior point on the left vallecula. | Vallecula Inferior Left | (VaInL) |
| 11 | Most inferior point on the right vallecula. | Vallecula Inferior Right | (VaInR) |
| 12 | Most anterior point of the anterior pharyngeal wall at the level of the velum tip as visualized in the midsagittal plane. | Velum Anterior | (VeAn) |
| 13 | Most posterior point of the posterior pharyngeal wall at the level of the velum tip, as visualized in the midsagittal plane. | Velum Posterior | (VePo) |
| 14 | Midpoint between VeAn and VePo (landmarks 12 and 13 as defined above). Midpoint was calculated based on VePo and VeAn landmark coordinates. | Midpoint between VeAn and VePo | (MidVe) |
| 15 | The most supero-posterior point of the velum. | Velum Back | (VeBa) |
| 16 | The inferior tip of the velum. | Velum End | (VeEnd) |
| 17 | The most anterior point of the pharynx at the level of the PNS in the axial plane as guided by the midsagittal plane of the pharynx. | Nasopharynx Anterior | (NpxAn) |
| 18 | Most posterior point of the pharyngeal wall at the level of the PNS in the axial plane as guided by the midsagittal plane of the pharynx. | Nasopharynx Posterior | (NpxPo) |
| 19 | Midpoint between NpxAn and NpxPo (landmarks 17 and 18). Midpoint was calculated based on NpxAn and NpxPo landmark coordinates). | Midpoint between NpxAn and NpxPo | (NpxMid) |
| 20 | Landmark placed on the soft tissue between the posterior vomer bone and the nasal crest of the palatine bone. | Posterior Nasal Septum | (NasalS) |
| **Reference Landmarks** |  |  |  |
| 21 | The most anterior point of the Anterior Nasal Spine. | Anterior Nasal Spine | (ANS) |
| 22 | The most posterior point of the Posterior Nasal Spine. | Posterior Nasal Spine | (PNS) |
| **Maxilla Landmarks** |  |  |  |
| 23 | The most postero-inferior point of the maxillary alveolar bone in the midsagittal plane (landmark location is between the first incisors) | Alveolar bone of incisor | (ABI) |
| 24 | The most posterior point of the incisive canal in the midsagittal plane of the maxilla. | Posterior edge of incisive canal | (PIC) |
| 25 | The intersection between the transverse palatine suture and the median palatine suture. | Palatine Sutures intersection | (PALS) |
| 26 | Midpoint between PIC and PALS (landmarks 24 and 25) along the median palatine suture on the maxilla. Coordinates calculated based on PIC and PALS landmark coordinates x, y, z. | Maxilla midpoint | (MMax) |

**Table 5.**
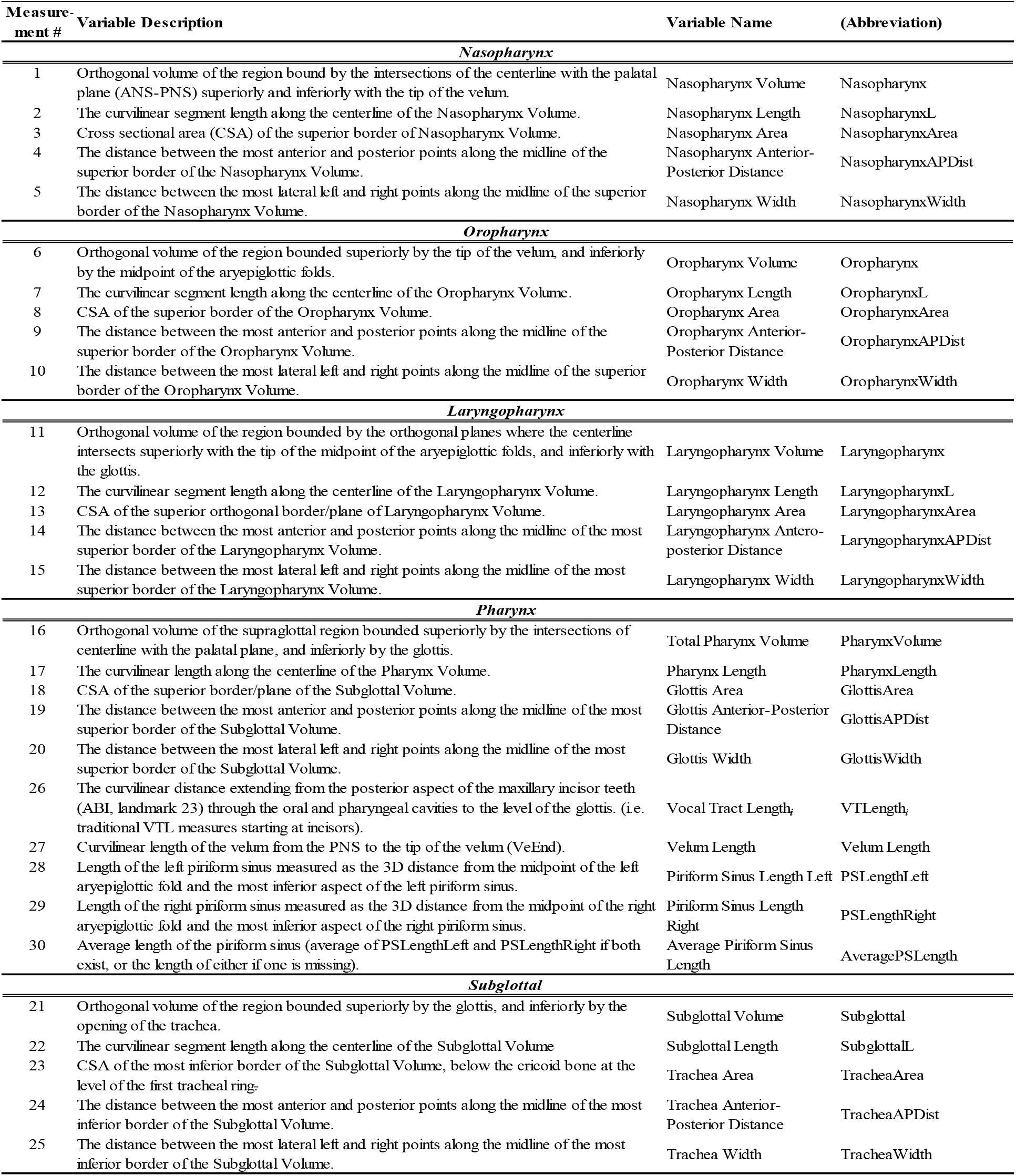
All upper airway variables examined. The 30 upper airway variables examined. Measurements extracted for each region include: the orthogonal volume, curvilinear/centerline volume-length, the orthogonal superior or inferior cross-sectional area (CSAs) of each of the subregions, as well as the anterior-posterior distance (APDist) and lateral width (Width) of each CSAs. All planar measurements are orthogonal to the centerline. The sum of the nasopharynx, oropharynx and laryngopharynx subregions was used to calculate pharynx volume and pharynx length (measurements 1-to-25). See text for additional vocal tract (VT) measurements (measurements 26-30).

### d. Pharynx centerline and data extraction protocol

A semi-automatic, centerline-based data extraction pipeline was developed to extract quantitative measurements from the 3D pharynx in MATLAB (The MathWorks, Natick, MA). First, the built-in marching-cube algorithm in MATLAB was used to generate 3D meshes of the pharynx model to serve as input to the pipeline [72]. This pipeline adapted the implicit fairing diffusion method to smooth the 3D pharynx meshes iteratively while preserving the intrinsic geometric properties [73]. Next, a level-contour-based centroid-extraction method was applied on the smoothed pharynx, obtaining a set of coordinates along the tubular center of the pharynx [74–76]. These coordinates were further interpolated and smoothed with the B-spline de Boor algorithm, generating a centerline representative of the center of the airway lumen [77, 78]. This centerline was then used as input to an in-house written script that calculated planes orthogonal (i.e., perpendicular) to the line segment formed by each centerline coordinate and its subsequent centerline coordinate. Finally, the intersections between the orthogonal planes and the 3D meshes were then extracted as boundary vertices. With the boundary vertices, cross sectional areas (CSAs) as well as additional variable measurements were calculated along the centerline. See Fig 2 for an illustration of the 3D pharynx model and the cross sections.

### e. Variable measurements

A total of 30 airway variables, as listed and defined in Table 5, were measured by the above-described protocol using planes orthogonal to the centerline. The variables extracted are described below and include overall pharyngeal length and volume, modified vocal tract length (VTLength*i*), velum length, and piriform sinuses length measurements, as well as measurements from the following four subregions: (i) Nasopharynx, (ii) Oropharynx, (iii) Laryngopharynx, and (iv) Subglottal. See Fig 1. Each subregion was isolated using its respective ‘landmark-derived planes’ orthogonal to the centerline using the following boundary definitions: The nasopharynx region was defined as an orthogonal volume bound by the intersection of the centerline with the palatal plane –formed by the anterior nasal spine (ANS) and posterior nasal spine (PNS) landmarks– superiorly, and with the tip of the velum inferiorly. The oropharynx region was defined as an orthogonal volume bound by the orthogonal planes at the tip of the velum superiorly, and by the aryepiglottic fold inferiorly. The laryngopharynx region, that includes the piriform sinuses, was defined as the orthogonal volume bound by the orthogonal planes at the midpoint of the aryepiglottic folds superiorly (the most curved point at approximate halfway between the base and tip of the aryepiglottic folds), and by the glottis inferiorly. The subglottal region was bound superiorly by the inferior boundary of the laryngopharynx region, and inferiorly by the first axial slice displaying the first tracheal ring. The first tracheal ring was used as a guide to the inferior border of the cricoid cartilage as the unossified cricoid cartilages in pediatric cases was difficult to delineate on the CT images [79].

Measurements extracted, as defined in Table 5, included for each region: the orthogonal volume, curvilinear/centerline volume-length, the orthogonal superior or inferior cross-sectional areas (CSAs, five total) of each of the subregions, as well as the anterior-posterior distance (APDist) and lateral width (Width) of each CSAs. The sum of the nasopharynx, oropharynx and laryngopharynx supraglottal subregions was used to calculate pharynx volume and pharynx length (Table 5, measurements 16 and 17). Additional vocal tract (VT) measurements (Table 5, measurements 26-to-30) included: Vocal Tract Lengthincisor (VTL*i*); calculated as the curvilinear distance extending from the posterior border of the maxillary incisor (seen as the most anterior landmark in Fig 1) to the glottis, representing the vocal tract portion of the upper airway starting at the incisor i.e. excluding the lip and teeth region. Velum length; calculated as the curvilinear distance extending from the PNS to the tip of the velum (VeEnd). Piriform sinus length (PSLength); left, right, and average PSLength, measured using the defined anatomical landmarks (PSSuL/R and PSInL/R).

### f. Statistical analysis

All statistical analyses were performed in R. A linear mixed-effect model was used to capture sex-specific growth in young children and allow for developmental comparison with adult pharyngeal morphology. This model, using the lmerTest package for mixed-effects in R, accounted for the repeat scans included in the dataset from individuals with multiple visits. The model was specified as follows:

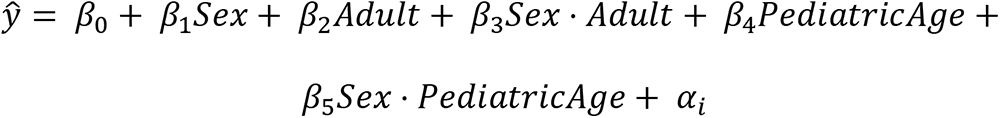

with “Adult” a dummy variable for adult subjects, “PediatricAge” the age for non-adult subjects (0 for adults), and α_*i*_ a random per-subject effect.

Outliers were first excluded using the model, by removing data points with residuals exceeding 2.576 of standard deviation, as described in [23, 71]. The model was then refitted on log-scale for each of the variables to assess for growth trends and sex differences.

Likelihood ratio test (LRT) was conducted to assess overall age effect in the first five years of life. To assess sex-differences, Wald test was performed at three time points: age-groups < 1 year, 5-years and adults. Tests were conducted at a nominal significance level of α = 0.05; in Table 6, significance at the stringent Bonferroni corrected level (<.0004) was also indicated.

**Table 6.**
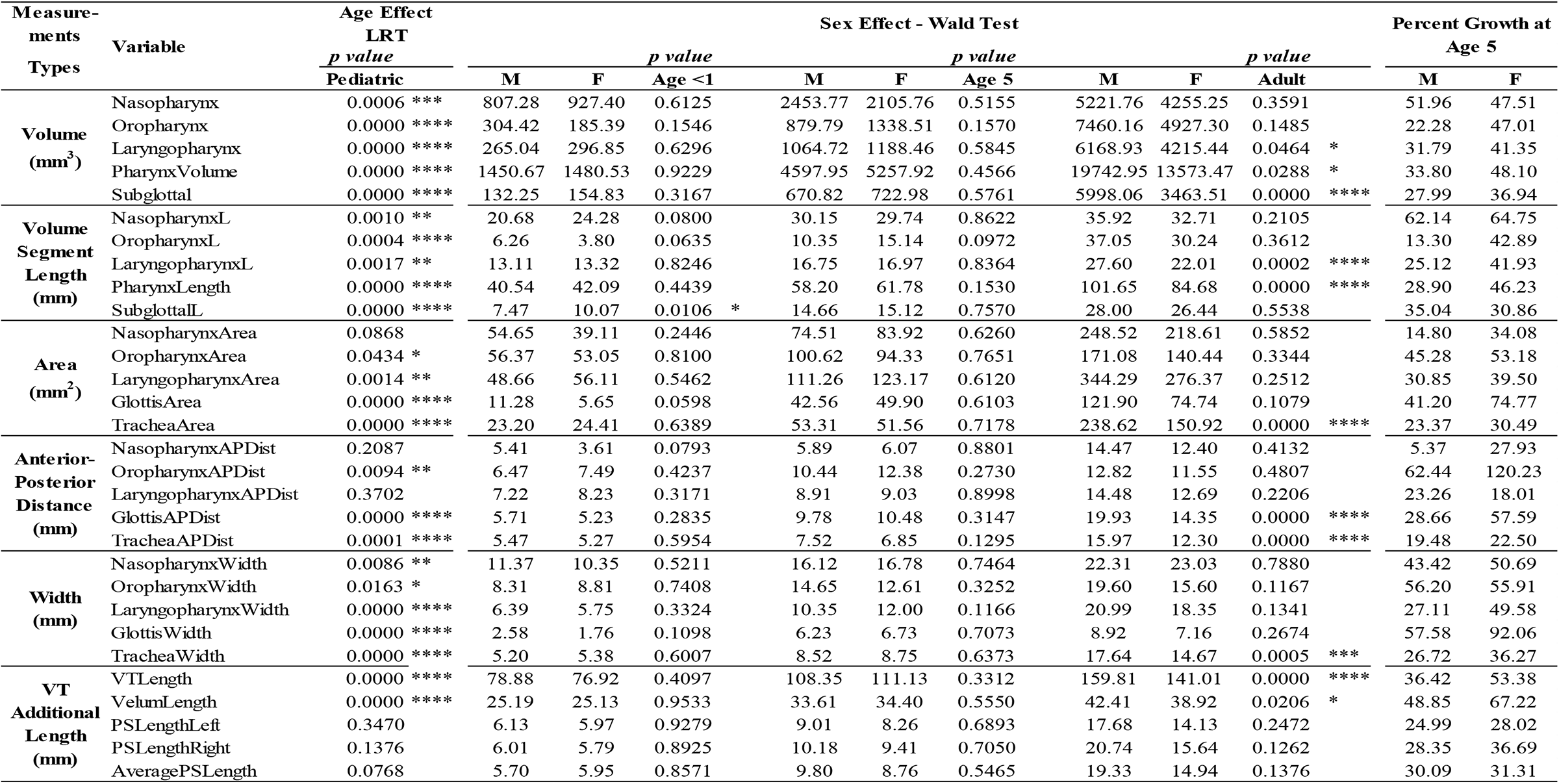
Likelihood ratio test and Wald test results. Likelihood ratio test (LRT) results for age effect, and sex-effect using the Wald test at age <1 year, 5 years, and adults. Significant differences are denoted with asterisk (* <.05; ** <.01; *** <.001; **** <.0004 Bonferroni corrected value). Also, percent of adult size at age 5 years (final column) for each of the 30 variables as listed by measurement type (column 1), and airway sub-regions (column 2). Percentages are based on the point estimate of modelled means, see Fig 3 for 95% confidence intervals. The regions are listed superior to inferior with supra-laryngeal (above glottis) measurements listed first. Refer to Fig 1 to visualize variables and subregions, and Table 5 for variable definitions.

Finally, using point estimate of modeled means, percent growth at age 5-years was calculated using data at age-group < 1 year, and adults for the purpose of gaining insight on upper airway growth type as described by Scammon [80]. Scammon determined two primary postnatal growth types, neural and general growth types, or their combination, to characterize growth of head and neck structures. Furthermore, he noted that while all primary growth types are characterized by a period of rapid growth during infancy, by early childhood neural growth type achieves greater than two-thirds of the adult size, while somatic growth type barely achieves a quarter of the adult size [80].

## Results

Measurements extracted for males and females are displayed in Figs 3A-C for each pharyngeal subregion (naso-oro-laryngo-pharynx), Fig 3D for the entire pharynx (supraglottal region), and Fig 3E for the subglottal region with sex-specific linear fits and confidence intervals at age <1 year, 5-years, and adults. The plots also include a second y-axis depicting the percent growth of adult size. Statistical analysis results are also summarized numerically in Table 6. Significance at the .05, < .01, < .001, and Bonferroni corrected < .0004 levels are marked with one, two, three and four asterisks respectively in Table 6.

**Fig 3.**
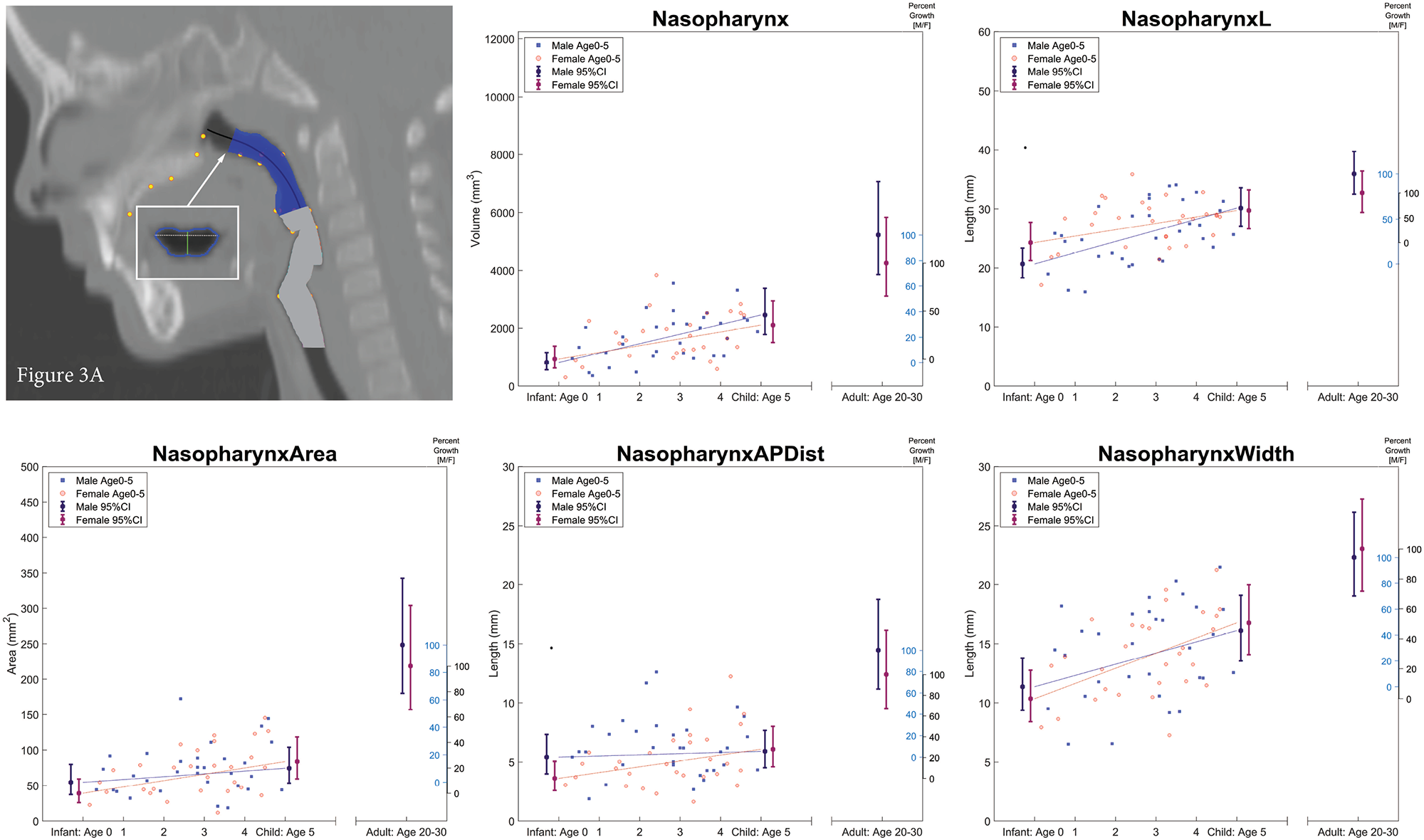
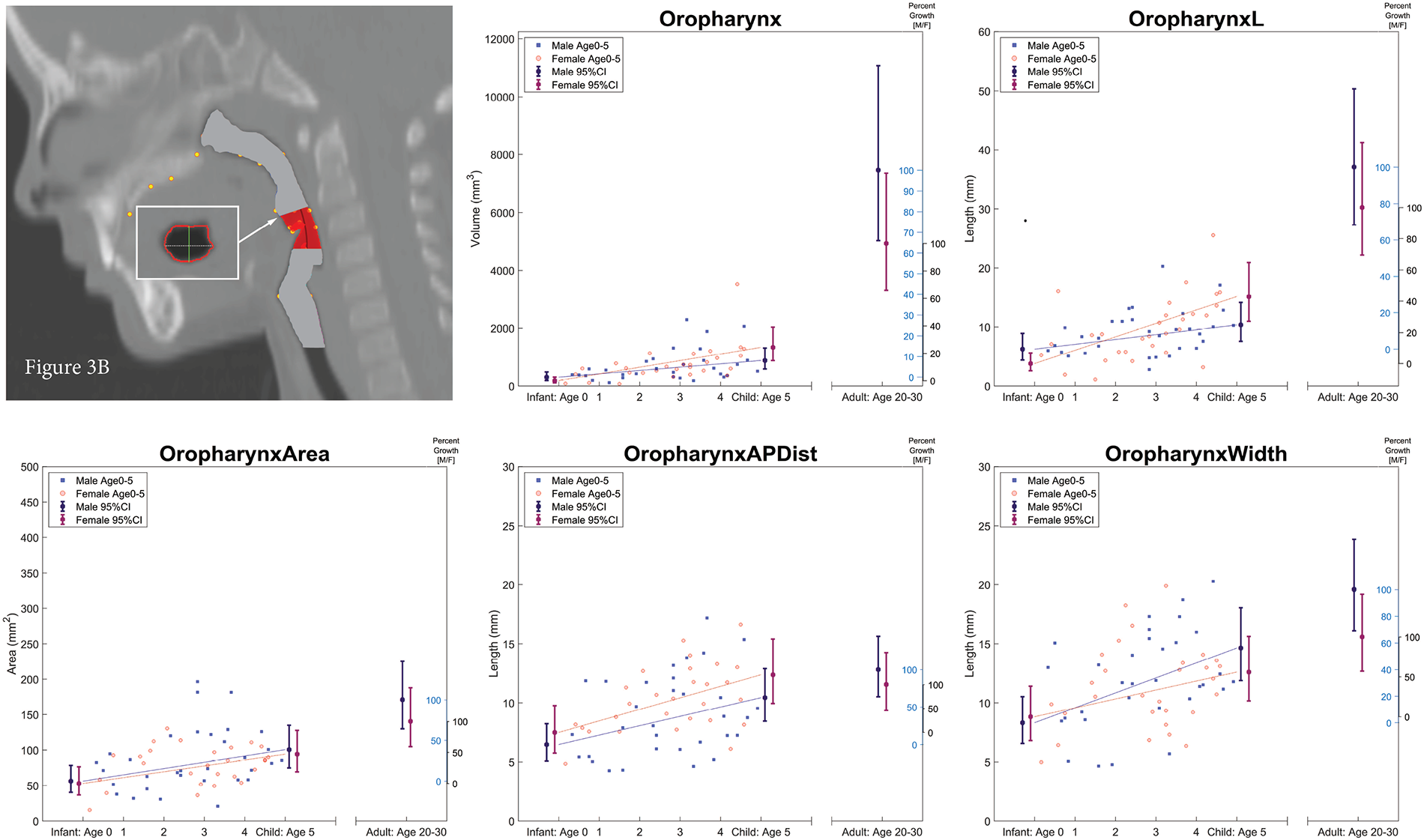
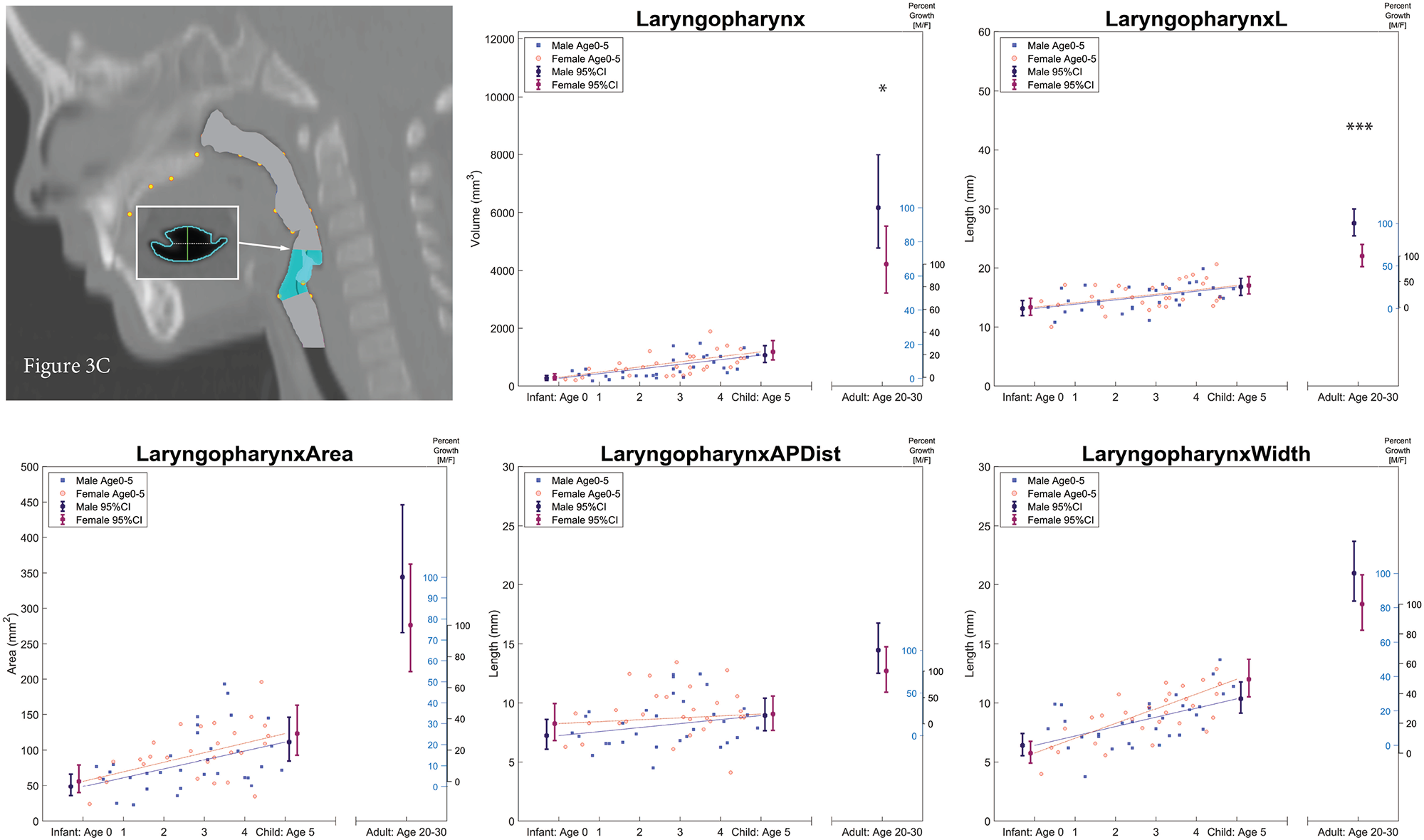
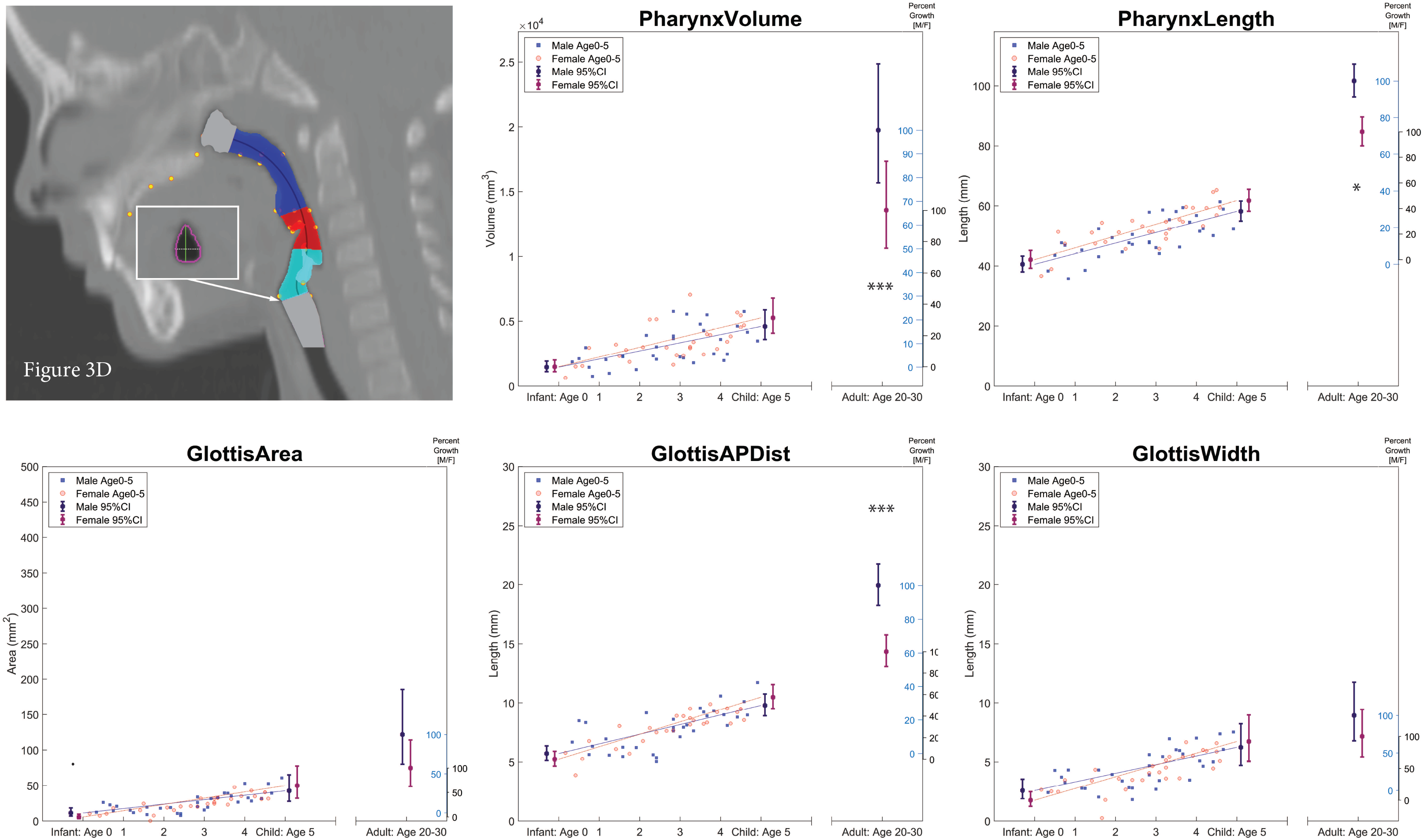
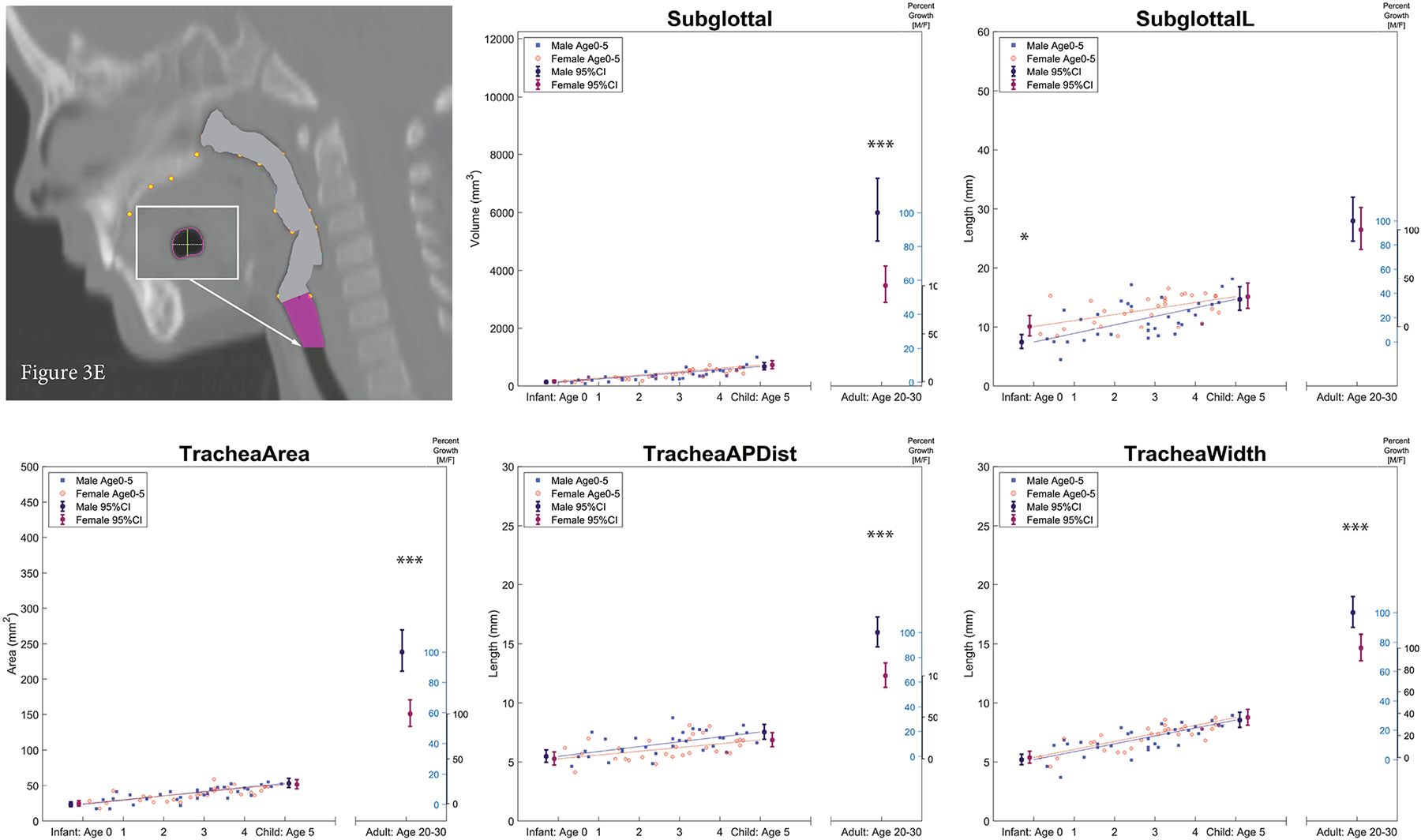
Measurements extracted for pharyngeal region (3A. Nasopharynx; 3B. Oropharynx; 3C. Laryngopharynx; 3D. Pharynx; and 3E. Subglottal). Measurements extracted for each pharyngeal region (3A to 3E) is depicted in top panel image on left, with measurements as defined in Table 5 and consisting of: Top panel; the orthogonal volume, and the curvilinear/centerline volume-length. Bottom panel; the orthogonal cross-sectional area (CSA) (superior 3A, or inferior 3B-3E), its anterior-posterior distance (APDist), and lateral width (Width). Plots include measurements for male in blue filled square symbols, and for female in red open circle symbols. Pediatric data include linear fits for males (blue solid line) and females (red dashed line). Point estimate of modeled means and confidence intervals are plotted for adult data, and at ages 0 and 5 years respectively for males (purple) and female (magenta). The second Y-axis reflects the percent growth for males (blue, inwards tick orientation) and females (black, outwards tick orientation). Significance for sex differences at birth, age five and/or adults are denoted with single, double, and triple asterisk(s) at the interval plots using the nominal α < 0.05, < 0.01, and < 0.001 levels respectively; the numeric p values are displayed in Table 6.

### a. Age Effect

As expected, likelihood ratio test results confirmed that all airway measurements for total and subregion volume, length (including VTL*i*), and width (lateral) exhibited statistically significant growth in size during the first five years of life (< 1 year to 5-years) (Table 6). Also, four of the five CSAs examined displayed significant growth in size, except for the CSA at the level of the nasopharynx (*p* = 0.0868). Similarly, the linear anterior-posterior distance (APDist) measurements displayed significant growth except for the two measurements at the level of the nasopharynx and the laryngopharynx (*p* = 0.2087 and 0.3702 respectively). Limited growth was noted for average piriform sinus length (AveragePSLength; *p* = 0.0768), but growth in Velum Length was highly significant (*p* < .0001).

Compared to the mature adult airway, both male and female pediatric upper airway dimensions by age 5-years were significantly smaller (with higher percent growth in females than males as discussed below). However, one exception was the oropharynx anterior-posterior distance (APDist) and to some extent oropharynx width, where by age 5, children had essentially attained their adult size (see Fig 3B).

### b. Sex Effect

The Wald test performed on pediatric data indicated that only subglottal length (SubglottalL) showed significance (*p* = .0106) at age <1 year with females’ mean length being longer than males (see Table 7 mean (s.d.); M=8.06 (3.02) mm; F=10.56 (3.17) mm). No statistical significance was detected for any other variable at age-group <1 year or age-group 5-years. However, sex differences – though not significant – were noted at age group < 1 year in the volume-length of the nasopharynx (*p* = 0.08) and the oropharynx (*p* = 0.0635) subregions with females having longer measurement than males; also, differences in nasopharynx APDist (*p* = 0.0793) and glottis area (*p* = 0.0598) with males having larger measurements than females. By age 5-years, insignificant differences in the volume-length of the oropharynx subregion was noted (*p* = 0.0972) with females having slightly longer measurements than males (see Table 7; M=10.43 (3.57) mm; F=14.28 (7.21) mm).

**Table 7.**
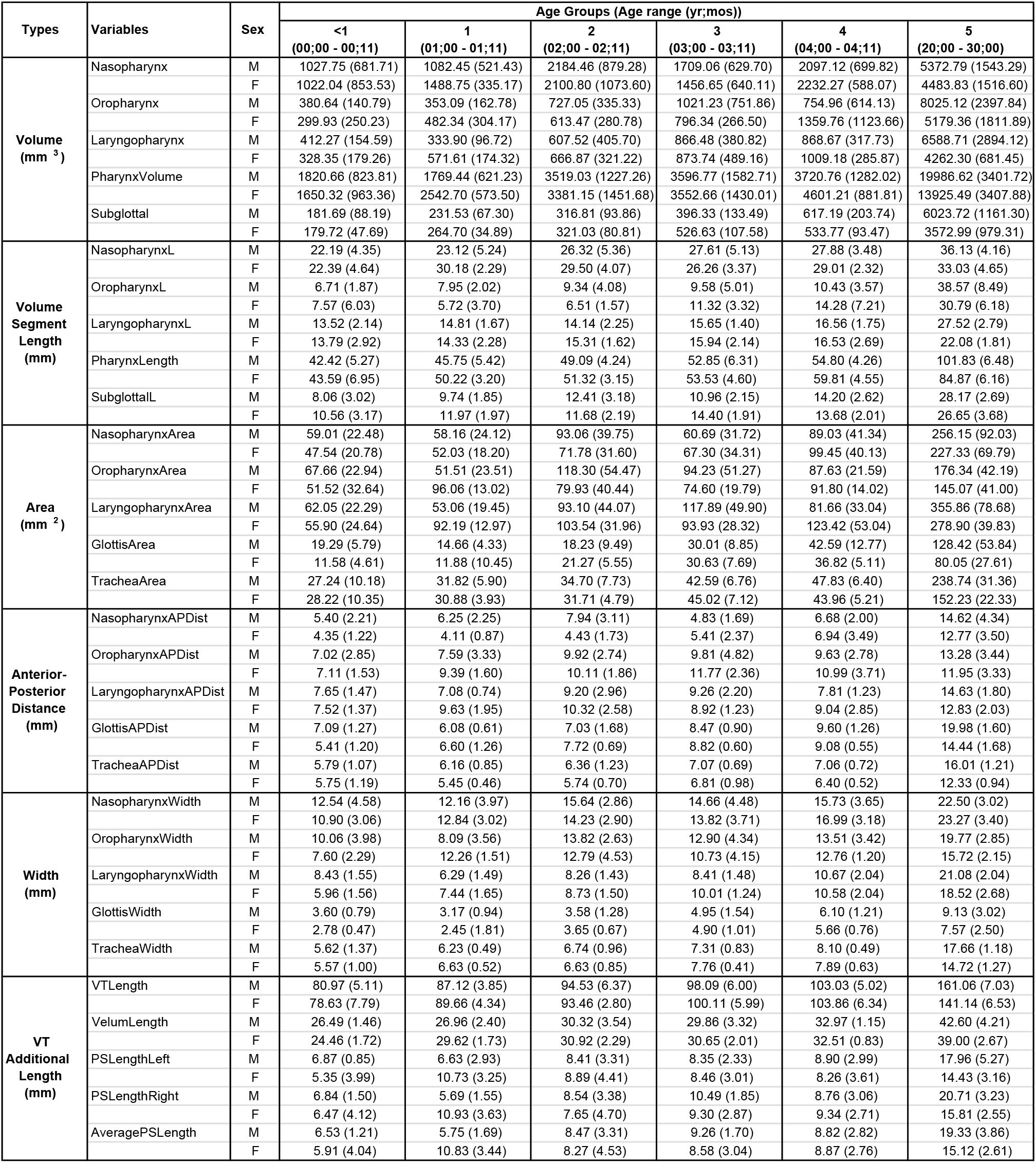
Age-specific mean (standard deviation) of the different measurement types for each pharyngeal region. Age-specific mean (standard deviation) of the different measurement types for each of the variables examined with M/F denoting the average Male/Female measurements. Age groups as specified in Table 3.

As for adults, sexual dimorphism was highly significant for overall pharynx length and VTL*i* (*p* < .0001; M=161.06 (7.03) mm; F=141.14 (6.53) mm), and significant for pharynx volume (*p* < .05; M= 53.73 (16.43) cm^3^; F= (44.84 (15.17) cm^3^), with males having larger measurements than females (see Tables 6 and 7). Sexual dimorphism was also present in laryngopharynx length (*p* < .0004) and volume (*p* < .05), as well as subglottal volume (*p* <.0004), with highly significant differences in Glottis APDist (*p* < .0004) superiorly and CSA at tracheal ring inferiorly (*p* <.0004; including differences in tracheal APDist (*p* <.0004) and width *p*=.0005) with males having larger measurements than females.

### c. Percent Growth

As displayed in second y-axes of Fig 3 plots, and listed in Table 6 (final column), percent growth assessment based on modeled point estimates, revealed that by age five-years, female upper airway measurements were closer than male measurements to the adult mature size in 26 out of the 30 upper airway variables examined. (See the tabulated average data per age-group in Table 7.) Female data revealed 9 out of 30 variables to have reached over 50% of adult size nasopharynx volume-length (64.75%), nasopharynx width (50.69%), oropharynx area (53.18%), oropharynx APDist (120.23%), oropharynx width (55.91%), glottis area (74.77%), glottis APDist (57.59%), glottis width (92.05%), and VTL*i* (53.38%). In contrast, males had only 5 of the 30 variables reach over 50% of the adult mature size: nasopharynx volume (51.96%), nasopharynx volume-length (62.14%), oropharynx APDist (62.44%), oropharynx width (56.20%), and glottis width (57.58%). The only 4 measurements where both male and female growth reached over 50% of their respective adult size were: nasopharynx length, oropharynx APDist, oropharynx width, and glottis width.

## Discussion

This study addresses a void in normative data on the upper airway during the first five-years of life. After developing a protocol that controls for variables that can affect measurement accuracy (e.g., head position, sedation, threshold for airway segmentation), CT studies from 61 typically developing pediatric and 17 adults were used to quantify the multidimensional growth of the airway systematically with respect to age and sex. Our findings are novel in that, to our knowledge, this is the first study that examines the birth-to 5 years age range, as compared to adults, using a comprehensive set of 2D and 3D measurements from the choanae to below the cricoid ring (opening to the trachea), including: supra- and sub-glottal volume and length, naso-oro-laryngo-pharynx subregion volume and length, each subregion’s superior and inferior CSA, and their antero-posterior and transverse/width distances. Additionally, the data were collected using a protocol that included a well-defined and established threshold for airway segmentation, and a semi-automatic centerline that we developed for the extraction of accurate measurements to quantify the upper airway using the natural anatomic orientation of airflow for respiration and speech production.

The use of a centerline, as an added methodological consideration, is critical for obtaining accurate measurements of the airway. As summarized in Table 1, two studies [35, 49] have used a centerline to quantify the airway, but Chiang et al. [35] is the only study to date that performed a centerline-based technique to obtain quantitative data on the growth and development of the airway. However, their measurements stopped at the level of the epiglottis, and they did not include the pre-pubertal age group.

The upper airway subregion dimensions are sensitive to altered head and tongue posture, particularly for 3D assessment but also for 2D measurements as Gurani et al. [54] point out.

Given the need for a valid method to classify head position of imaging studies, our laboratory first developed a reliable protocol that uses 14 landmarks to account for both head and neck positions [68]. We therefore first employed this protocol for the selection of cases with a neutral head position for inclusion in this study, then applied the centerline protocol.

Given all the methodological considerations we accounted for, the attrition rate of cases included in this study from the imaging studies available in our database was high. We retained only 19% of the cases reviewed. Given this rigorous approach to control for positioning and other potential confounder, we expect our findings to reliably reflect typical airway growth. Furthermore, we anticipate that the inclusion of additional cases in future studies, using the above-described airway data extraction protocol, will further strengthen present findings and observations.

### a. Age Differences

Our findings reflect persistent positive increase in size for all variables examined during the first five years of life for all measurements in all subregions as displayed in Fig 3, with the means per age group summarized in Table 7. The age effect of the likelihood ratio test confirmed the significant growth in size for 24 out of the 30 variables studied with some variables displaying more rapid and extensive growth than others. Such findings provide insight on the proportional and relational growth of upper airway dimensions with age during anatomic restructuring (e.g., hyo-laryngeal descent).

As expected, all pediatric airway dimensions were substantially smaller than adult dimensions except oropharynx APDist and to some extent oropharynx width (see Fig 3). Abramson et al. [20] also found that volume, CSA and transverse measurements –but not AP dimensions– of the pediatric naso-oropharynx airway were significantly smaller than adult measurements. This rapid and early maturation in oropharynx APDist dimension corresponds to our previous research findings where growth of oral structures in the horizontal plane, in line with neural growth, achieved most of their growth towards the adults size by age five-years [23, 66, 71]. Alternatively, though unlikely, it is possible that hypertrophy of lingual tonsils interfered with lumen APDist measurements. As for growth in width, while not as fully developed as AP dimensions by age 5, oropharynx lateral dimensions had reached over 55% of the adult, suggesting that growth in this subregion undergoes a combination of neural and general growth types, which Scammon had noted is present in the growth of structures in the neck region.

Volume, volume-length and width measurements increased with age for all pharyngeal subregions, consistent with prior studies on upper airway development in infant and pre-pubertal children [20, 24]. The CSA measurements in this study were extracted from anatomical landmarks representing the superior and inferior borders of the pharyngeal subregions. This is in contrast to the typical approach of measuring minimum or maximum CSA to examine sites of constriction for assessment of patients’ risk for OSAS [39, 40, 42, 43, 45, 49, 81]. Furthermore, we used oblique planes – orthogonal to the centerline – which cannot be compared to studies that used the axial plane, as in most of the above listed studies.

A factor that further complicates comparisons, including within-study cross-sectional comparisons, is the hypertrophy of tonsils in young children that follow a lymphoid growth type. In particular, nasopharyngeal tonsils referred to as adenoids, where hypertrophy is the highest in 4-6-year old children [82]. Keeping these issues in mind, among the five CSA measurements in this study, the nasopharynx region was the only site that did not have a significant age-effect.

Similarly, the APDist measurements in the nasopharynx and laryngopharynx were the only sites that did not have significant age effect between the ages < 1-year and 5-years. Such findings could be attributed to adenoid hypertrophy, typically occurring between the ages 2-6 years, that diminish airway dimensions [83–85]. The decrease of mean nasopharynx CSA and APDist measurements per age group can be noted in Table 7, with changes most evident between the ages 2-to-4 years in this study.

In all age groups, the APDist dimensions at the nasopharynx, oropharynx and trachea (Fig 1a, b, e) were smaller than the width/transverse measurements, but larger than width/transverse measurements at the glottis (Fig 1d, and Table 7). Such findings are in line with Abramson et al. [20] who reported significant upper airway growth along the transverse dimension with age, where the airway becomes more elliptical in shape. Similarly, Machata et al. [26] using MRI studies of children ages 0-6 years, reported smaller anteroposterior dimensions than transverse dimensions for all ages at the level of the soft palate, the base of the tongue, and the tip of the epiglottis. In contrast, the laryngopharynx APDist dimensions (Fig 1c), were larger than width/transverse measurements from birth to age 3, but became smaller than width/transverse measurements at age 3 and beyond, which likely contributed to the absence of age effect for APDist.

Changes in APDist versus width dimensions could be attributed to the cartilaginous composition of the larynx. The laryngopharyngeal cross section in this study was designed to capture its surrounding structures - the aryepiglottic folds on each side, the laryngeal vestibule anteriorly, and posteriorly by arytenoid cartilages, corniculate cartilages and the interarytenoid fold. Cartilage ossification is usually not observed until past age 20 years, and the pediatric laryngopharynx region is often described to be “featureless” and difficult to assess using commonly acquired medical images [79]. We employed an established method for airway segmentation that uses image-specific airway thresholds, and therefore are confident that our data is reflective of airway development. Since we used landmarks on the aryepiglottic folds that connect to the piriform sinuses on each side of the cavity, aditus of larynx, the laryngopharynx width/transverse measurement in this study excluded the piriform sinuses (see Fig 1c). The significant age effect along this transverse dimension is therefore truly reflective of the strong lateral growth in the laryngopharyngeal region.

The CSA and transverse width measurements at the level of the glottis are smaller than the area and width at the first tracheal ring immediately inferior to the cricoid (i.e., subglottal region); however, the average APDist measurement of the glottis is larger than the APDist dimension at the level of the first tracheal ring at ages <1-year, 5-years, and in adults. This finding is consistent with Luscan et al. [86], who concluded that “the cricoid has a round shape regardless of the child’s age.” Indeed, the mean APDist and width measurements were very close or similar for all ages at the level of the first tracheal ring proximal to the inferior border of the cricoid, and comparable to the cricoid outlet’s (to trachea) anteroposterior and transverse interior diameters of Liu et al. [87]. While growth trends were comparable, our measurements were closer to those of Liu et al [87] than to those of Luscan et al. [86], and indicate the importance of methodological considerations, including having well-defined data extraction protocols such as the determination of an appropriate threshold level (HU) to segment the airway.

Growth in VTL*i* was significant during the first five years of life, confirming an increase of about 3 cm, which is consistent with VTL findings to date [23] and reflects that this modified measure captured growth in both the oral and pharyngeal portions of the VT. The measurements in this study were smaller than what has been reported previously, which is to be expected given the modified length measure had an onset at the posterior margin of the incisors in lieu of the typical anterior margin of the lips. Findings of a significant age effect on velum length were comparable to values reported by Perry et al. [88] and Yi et al. [27]. Closer examination of the developmental data on pharyngeal length and pharyngeal volume revealed a close relationship particularly after about age 2. Before age 2, the growth rate was slightly more pronounced in length than in volume, likely due to the drastic anatomic restructuring of the skeletal framework in the region of the pharyngeal cavity, including hyo-laryngeal descent, and rapid neural growth in length of the second cervical spine (C2) [68].

As for the piriform sinuses, our findings revealed a borderline average PSLength age effect (*p*=.077) with average measurements per age-group summarized in Table 7. The pediatric average PSLength measurements ranged from 6 to 9 mm at ages <1-year to 5-years; whereas the adult average PSLength measurements ranged from 1.5 to 1.9 cm. While the development of piriform sinuses has not been examined to date, and therefore comparative measurements were not available, adult PSLength measurements were comparable to the 1.6 to 2 cm piriform sinus depth measurements of Dang and Honda [89–91]. This similarity was despite the fact that our PSLength measurement extended from the most inferior aspect of the piriform sinuses to the midpoint of the aryepiglottic folds, which is beyond the arytenoid apex plane used by Dang and Honda [89]. This could be in part due to differences in imaging modality used (CT vs MRI) and/or segmentation thresholding levels used to obtain reliable airway measures, particularly given the small size of this region of interest. Additional factors include methodological differences (oblique vs, axial plane) in obtaining measurements, as well as the height of participants, which has been shown to be related to vocal tract length [24] and pharyngeal dimensions [47]. The piriform sinuses play an important role during swallowing by diverting liquids around the aditus of the larynx and into the esophagus. They also affect speech acoustics and attenuate the vocal tract resonant frequencies in adults [89, 92–94] by an estimated range of 5% of formant frequencies [89]. Thus, detailed developmental data on the piriform sinuses would provide needed normative data and could help provide insight on pediatric dysphagia.

Furthermore, such data can be used to implement modeling [95] to systematically examine the effect of the piriform sinuses on the resonances of the developing vocal tract, particularly given the intriguing findings that formant frequencies reportedly remain stable during the first 24-to-36 months of life [14, 15, 96], despite documented increases in vocal tract length [2, 23, 24].

In summary, the upper airway dimensions reveal persistent growth during the first five years of life, with some dimensions growing at a faster pace than others. Growth in the vertical and transverse/lateral dimensions are more pronounced than growth in the AP dimension.

### b. Sex Differences

As depicted in Table 6, sexual dimorphism was present in a number of supra- and sub-glottal variables in adults. However, while there was evidence towards sexual dimorphism in all three supra-glottal regions for a number of variables at age <1 year (specifically, larger nasopharynxL and oropharynxL in females; also, larger nasopharynx APDist and glottis area in males), with the larger oropharynxL persisting in females at age 5 years, none of these supra-glottal or pharyngeal variables were significant in children.

As for the subglottal region, only subglottal volume-length displayed significant sexual dimorphism at age < 1-year (with males shorter than females), but not at age 5-years or in adults (see Table 6). To our knowledge, this specific subglottal volume-length measurement has not been examined, despite its importance in procedures like tracheotomy [97], and laryngotracheal infections/diseases including SIDS [98] where the incidence is higher in males [99]. However, two studies have performed distance measurement in this region, specifically anterior commissure to first tracheal ring [100], and vocal folds to the cricoid cartilage [101]. Contrary to present findings, Khadivi and colleagues, who used laryngoscopy to collect subglottal length data from 82 adults (57 males and 25 females), documented significant sexual dimorphism.

While our measurements for pediatric subglottal length are comparable to the normative values reported by Sirisopana and colleagues, from the CT scans of 56 children (29 males, 27 females), they unfortunately neither assessed for sex differences, nor reported sex-specific measurements given their primary focus on tracheal tube design.

Despite methodological differences, the absence of sexual dimorphism in pediatric upper airway data for most of the measurements analyzed in this study was consistent with past studies [22, 24, 29]. Barbier et al. [29] did not find sex difference in pre-pubertal data but suggested that sexual dimorphism in VTL emerged during puberty. Among studies reporting regional upper airway normative data for the age range between 0-to-5-years, Abramson et al. [20] found no difference in naso-pharyngeal airway size or shape between the sexes in children, but reported longer airway length in post-pubertal males. Jeans et al. [83], using lateral cephalometric radiographs to study the nasopharyngeal airway area – comparable to our nasopharyngeal region –, however, reported mild decreases in nasopharyngeal area in both 3-to-5-year old males and 3-to-6-year old females. Sex-differences in the same region using an anteroposterior distance measure in the midsagittal plane of medical imaging studies (MRI & CT), referred to as oropharyngeal-width, have similarly been noted to display evidence, albeit not significant, towards sexual dimorphism in 3-to-4-year old children with males having larger width measurements [71]. Linder-Aronson et al. [85] noted that nasopharynx airway depth/AP dimension in males were consistently larger than females throughout ages 3-to-16 years. Sexual dimorphism in the pharyngeal portion of the VT in ages 8-to-19 years, has also been reported by Vorperian et al. [71], with the vertical nasopharyngeal length being longer in females than males and the vertical posterior cavity length being longer in males than females. In contrast, Yi et al. [27] reported no sex differences in any of their linear dimensions using MRI in infants and children up to 72 months. Rommel et al. [34], who used curvilinear length drawn on 2D X-ray images to assess naso-oropharynx segments, found no sex difference in children as young as 0-to-4-years. Griscom [102] similarly found no significant sex differences in trachea dimensions until late in adolescence. Definitive prepubertal sexual dimorphism of the pharynx thus cannot be confirmed with studies available to date.

Detecting sex differences is a difficult task given the critical methodological considerations outlined in our methods section and the importance of having a large number of participants per age-group. Statistical analysis methods can overcome differences in growth rate between males and females, such as implementing continuous-window comparisons across age [e.g.,71]. This latter approach was particularly effective in unveiling sexual dimorphism that does not persist during the course of development, since growth in females outpaces males during early development, but growth in males begins to outpace females during the peripubertal period, with sexual dimorphism emerging during puberty.

Sexual dimorphism in adults, however, was mostly present and aligned with research findings to date in pharynx volume, pharynx length [47], VTL [71], velum length [103], glottis APDist [47], subglottal volume [102], and tracheal dimensions [86]. The finding that subglottal volume-length, the only measurement that displayed significant sexual dimorphism at age < 1-year-old was not sexually dimorphic at age 5-years is not surprising, given growth rate differences in males versus females as noted above. However, the absence of differences in adults is likely due to both methodological differences and the limited number of adults in this study (*n*=17), given our stringent inclusion criteria.

Although present findings revealed significant prepubertal sex-differences only in the subglottal region, findings in this study and others as noted above, in the naso-oro-pharyngeal region, provide sufficient justification to further examine this issue using a larger number of cases particularly given the above noted critical methodological considerations, and the nuance of growth rate differences between the two sexes. Such a conclusion is further supported by auditory-perceptual and acoustic findings where Bloom et al. [104] reported that adults accurately identified 3-month old infants’ vocalizations as boy vs girl. The only acoustic difference was the feature of nasality with girls’ vocalizations being more nasal than boys.

Furthermore, several studies have reported sex differences in vowel formants (i.e., vocal tract resonant frequencies) in children as young as 3 or 4 years of age [105–107].

### c. Percent Growth

The growth pattern of anatomical structures in the craniofacial and upper airway region are known to be non-uniform with the primary growth types in this region being neural, general/somatic, and lymphoid. Since we did not include adenoid or tonsil measurements, we will limit this discussion to the first two types. Both neural and general growth types display rapid growth during the first few years of life. Scammon [80] summarized schematically, however, that by age 5, the percent of the adult mature size reached was drastically different for the neural (∼80%) versus the general (< 40%) growth types. He also noted that growth in the neck region could be a combination of both neural and general primary growth types [80]. The final column of Table 6 presents the model-based point-estimates of percent growth of the adult mature size by age five years for all thirty variables examined in this study.

The general finding that by age 5, female upper airway dimensions were larger than males is not surprising since typically females have a faster growth rate during childhood and reach the adult mature size sooner than males. Based on findings to date, structures in the upper airway were expected to follow a mostly somatic growth type or a composite growth of somatic and neural growth types [23, 108, 109]. Despite differences in VTL versus VTL*i* measurements (where the onset of the former at the anterior margin of the lips and the latter is at the posterior margin of the incisors), the general growth findings in this study are in line with the expected growth trend indicating that VTL*i* growth type is predominantly hybrid somatic/neural in females (53.38%) and somatic in males (36.42%) [23]. Similarly, pharynx volume, pharynx volume-length and all other pharyngeal subdivision volume and volume-length results except for the nasopharynx subregion confirmed the predominant somatic growth types at age 5-years, in line with the reported growth patterns for pharyngeal cavity length and VT vertical data [23]. As for the nasopharyngeal region, the expected hybrid somatic/neural or combination growth type, was indeed reflected in present findings where both male and female nasopharynx volume and volume-length measurements ranged between 47% and 65% of adult size by age 5-years. The oropharynx AP dimension exceeded 60% for males and 85% for females, suggestive of a more pronounced hybrid somatic/neural growth type for the males and neural growth type for the females. These findings are in line with Vorperian et al. [23] where structures in oral region in the horizontal plane reached maturation earlier than structures in pharyngeal region in the vertical plane. Percent growth analysis (in Table 6, last column) also reflect the presence of sex-specific differences in the combination of growth types for upper airway structures. Since multiple factors contribute to growth, it is possible that such sex-specific differences further contributed to the difficulty in detecting sexual dimorphism during early childhood.

## Conclusions, Study Limitations, and Future Direction for Research

This study, using CT studies, provides data quantifying the 3D growth of the upper airway with minimal methodological concerns for an age group with scant normative data. Findings confirm persistent growth of the upper airway during the first five years of life with growth in the vertical and transverse/lateral dimensions having a faster pace and greater prominence than growth of anteroposterior dimension. Findings also reveal that at age 5, females have larger airway dimensions than males. Such findings confirm the importance of studying sex-specific developmental changes of the upper airway in 3D. A better understanding of pharyngeal functions and disorders will require further, more detailed examination of the developmental changes in pharyngeal length versus volume and the piriform sinuses, as well as re-examination of prepubertal sex-differences.

Our painstaking efforts to optimize accurate and reliable upper airway measurements limited the sample size. Specifically, the attrition rate of only retaining retrospective imaging studies with a neutral head position was high given the head position protocol we applied [68]. Also, despite the imaging protocol at the University Hospital to maintain the head in midline, it is likely that some rotation was present. However, given that the airway is functional during imaging (breathing and swallowing), it is difficult to determine source of asymmetry as noted with the piriform sinuses, where we resorted to averaging them in this study. Furthermore, airway anatomic measurements can be affected by breathing phase where significant effect of breathing phase in the oro-laryngo-pharyngeal region has been reported [34]. Such concerns, including the above suggested assessments, could be addressed by replicating this study using a larger sample size, and ideally increasing the age range to cover the entire developmental period particularly ages 5-to-20 years.

Aside from using a larger sample, having relevant demographic information such as height, weight, and race would be valuable to include. Given the retrospective nature of this study, relevant demographic information was not available for all cases. Our imaging database however, was representative of regional Dane County demographics, with growth between the 10^th^ and 95^th^ percentiles. We believe our present findings are representative of typical growth since the natural variability in craniofacial dimensions within individual races is related to the natural variability or variations within the different racial/ethnic groups [110].

Aside from the use of 3D anatomic landmarks and establishing standardized procedures to minimize, if not eliminate, methodological limitations on measurement accuracy and reliability, it is also imperative to establish standardized definitions of the pharyngeal subregions using well-defined anatomical boundaries. This will facilitate accurate representation and comparison of the developmental morphology of the aerodigestive and vocal tract, within and across disciplines, including comparison between different imaging modalities. This will undoubtedly enrich our understanding of the growth of a region that serves multiple life-functions since each modality has its strengths and limitations. For example, while MRI provides more accurate information on the growth of lymphoid tissue in the pharyngeal region, CBCT could address concerns on the effect of body position and gravity on soft-tissue structures for obtaining reliable airway dimensions. The feasibility of comparing developmental findings across disciplines, including the individual and relational growth of structures that provide the skeletal framework of the aerodigestive and vocal tract, will facilitate the understanding of upper airway pathophysiology, improve surgical planning such as estimation of laryngeal mask airway size or endotracheal tube diameter, evaluation of pharyngeal collapsibility in early childhood in the assessment of OSAS, and other upper airway anomalies including swallowing difficulties and speech disorders. Such information would also facilitate the advancement of developmental models to assess various typical and atypical functions related to airflow, swallowing, and speech production.

## Acknowledgments

We gratefully acknowledge Lindell R. Gentry, MD for his assistance in establishing the Vocal Tract Development Laboratory’s imaging database. We thank Gabe Jardim for establishing a protocol to obtain threshold for airway segmentation, and Drs. Andy Alexander and Michael Speidel for their guidance in resolving and confirming the reliability of this protocol. We also thank Drs. Moo K. Chung and Nagesh Adluru for their input on centerline development, Ben M. Doherty for his guidance with the data extraction script, Sophie D. Blankenheim for assistance with literature review and preparation of Table 1, and Dr. Jacqueline J. Houtman for comments on an earlier version of this manuscript.

## Supporting Information

**S1 Data. Data used in this study**. Variable measurements as described in Table 5.

## References

1. Marcus CL, Smith RJH, Mankarious LA, Arens R, Mitchell GS, Elluru RG, et al., editors. Developmental aspects of the upper airway: Report from an NHLBI Workshop, March 5-6, 2009. Proceedings of the American Thoracic Society; 2009.

2. Vorperian HK, Kent RD, Lindstrom MJ, Kalina CM, Gentry LR, Yandell BS. Development of vocal tract length during early childhood: A magnetic resonance imaging study. The Journal of the Acoustical Society of America. 2005;117:338–50. doi: 10.1121/1.1835958.

3. Laitman JT, Crelin ES. Postnatal development of the basicranium and vocal tract region in man. In: Bosma J, editor. Symposium on the development of the basicranium. Bethesda, MD: DHEW Publicaiton No. 76-989, PHS-NIH; 1976. p. 206–20.

4. Roche AF, Barkla DH. The level of the larynx during childhood. 1987:645–54.

5. Enlow DH, Hans MG. Essentials of Facial Growth. Philadelphia: W.B. Saunders Company; 1996 Jun 1996. 303 p.

6. Moss ML. The functional matrix hypothesis revisited. 1. The role of mechanotransduction. American Journal of Orthodontics and Dentofacial Orthopedics. 1997;112(1):8–11. doi: https://doi.org/10.1016/S0889-5406(97)70267-1.

7. Moss ML. The functional matrix hypothesis revisited. 2. The role of an osseous connected cellular network. American Journal of Orthodontics and Dentofacial Orthopedics. 1997;112(2):221–6. doi: https://doi.org/10.1016/S0889-5406(97)70249-X.

8. Moss ML. The functional matrix hypothesis revisited. 3. The genomic thesis. American Journal of Orthodontics and Dentofacial Orthopedics. 1997;112(3):338–42. doi: https://doi.org/10.1016/S0889-5406(97)70265-8.

9. Moss ML. The functional matrix hypothesis revisited. 4. The epigenetic antithesis and the resolving synthesis. American Journal of Orthodontics and Dentofacial Orthopedics. 1997;112(112):4–410. doi: https://doi.org/10.1016/S0889-5406(97)70049-0.

10. Carlson DS. Theories of Cranofacial Growth in the Postgenomic Era. Seminars in Orthodontics. 2005;11:172–83. Epub 2005.

11. Standerwick RG, Roberts WE. The aponeurotic tension model of craniofacial growth in man. Open Dent J. 2009;3:100–13. Epub 2009/07/03. doi: 10.2174/1874210600903010100. PubMed PMID: 19572022; PubMed Central PMCID: PMC2703201.

12. Castaldo G, Cerritelli F. Craniofacial growth: evolving paradigms. Cranio. 2015;33(1):23–31. Epub 2014/12/31. doi: 10.1179/0886963414Z.00000000042. PubMed PMID: 25547141.

13. Lieberman DE. Epigenetic Integration, Complexity, and Evolvability of the Head. In: Hallgrímsson B, Hall BK, editors. Epigenetics: Linking Genotype and Phenotype in Development and Evolution: University of California Press; 2011. p. 271–89.

14. Gilbert HR, Robb MP, Chen Y. Formant frequency development: 15 to 36 months. Journal of Voice. 1997;11(3):260–6.

15. Kent RD, Murray AD. Acoustic Features of Infant Vocalic Utterances at 3,6, and 9 Months. Journal of the Acoustical Society of America. 1982;72(72):2–353. doi: Doi 10.1121/1.388089. PubMed PMID: WOS:A1982PB08400008.

16. Bunton K, Leddy M. An evaluation of articulatory working space area in vowel production of adults with Down syndrome. Clin Linguist Phon. 2011;25(4):321–34. Epub 2010/11/26. doi: 10.3109/02699206.2010.535647. PubMed PMID: 21091205.

17. Kent RD, Vorperian HK. Speech Impairment in Down Syndrome: A Review. Journal of Speech, Language, and Hearing Research. 2013;56(1):178–210. PubMed Central PMCID: PMCNIHMSID: NIHMS 392496.

18. Eslami E, Katz ES, Baghdady M, Abramovitch K, Masoud MI. Are three-dimensional airway evaluations obtained through computed and cone-beam computed tomography scans predictable from lateral cephalograms? A systematic review of evidence. Angle Orthod. 2017;87(1):159–67. Epub 2016/11/02. doi: 10.2319/032516-243.1. PubMed PMID: 27463700.

19. Ono T, Otsuka R, Kuroda T, Honda E, Sasaki T. Effects of head and body position on two- and three-dimensional configurations of the upper airway. Journal of Dental Research. 2000;79:1879–84. doi: 10.1177/00220345000790111101.

20. Abramson Z, Susarla S, Troulis M, Kaban L. Age-related changes of the upper airway assessed by 3-dimensional computed tomography. Journal of Craniofacial Surgery. 2009;20:657–63. doi: 10.1097/SCS.0b013e318193d521.

21. Smitthimedhin A, Whitehead M, Bigdeli M, Gustavo N, Geovanny P, Hansel O. MRI determination of volumes for the upper airway and pharyngeal lymphoid tissue in preterm and term infants. Clinical Imaging. 2018;50:51–6. doi: 10.1016/j.clinimag.2017.12.010.

22. Ronen O, Malhotra A, Pillar G. Influence of Gender and Age on Upper-Airway Length During Development. Pediatrics. 2007;120:e1028–e34.

23. Vorperian HK, Wang S, Chung MK, Schimek EM, Durtschi RB, Kent RD, et al. Anatomic development of the oral and pharyngeal portions of the vocal tract: An imaging study. The Journal of the Acoustical Society of America. 2009;125:1666–78. doi: 10.1121/1.3075589.

24. Fitch WT, Giedd J. Morphology and development of the human vocal tract: A study using magnetic resonance imaging. The Journal of the Acoustical Society of America. 1999;106:1511–22. doi: 10.1121/1.427148. PubMed PMID: 10489707.

25. Litman RS, Weissend EE, Shrier DA, Ward DS, editors. Morphologic Changes in the Upper Airway of Children during Awakening from Propofol Administration. Anesthesiology; 2002.

26. Machata AM, Kabon B, Willschke H, Prayer D, Marhofer P. Upper airway size and configuration during propofol-based sedation for magnetic resonance imaging: An analysis of 138 infants and children. Paediatric Anaesthesia. 2010;20:994–1000. doi: 10.1111/j.1460-9592.2010.03419.x.

27. Yi X, Yao L, Yuan X, Wei Y, Wang Z. Linear dimensions of normal upper airway structure by magnetic resonance imaging in Chinese Han infants and preschool children. Sleep Medicine. 2017;37:98–104. doi: 10.1016/j.sleep.2017.06.011.

28. Fregosi RF, Quan SF, Kaemingk KL, Morgan WJ, Goodwin JL, Cabrera R, et al. Sleep-disordered breathing, pharyngeal size and soft tissue anatomy in children. Journal of Applied Physiology. 2003;95:2030–8. doi: 10.1152/japplphysiol.00293.2003.

29. Barbier G, Boë LJ, Captier G, Laboissière R. Human vocal tract growth: A longitudinal study of the development of various anatomical structures. Proceedings of the Annual Conference of the International Speech Communication Association, INTERSPEECH. 2015;2015-Janua:364-8.

30. Goncalves RdC, Raveli DB, Pinto Ados S. Effects of age and gender on upper airway, lower airway and upper lip growth. Braz Oral Res. 2011;25(3):241–7. Epub 2011/06/15. doi: 10.1590/s1806-83242011000300009. PubMed PMID: 21670855.

31. Sheng CM, Lin LH, Su Y, Tsai HH. Developmental changes in pharyngeal airway depth and hyoid bone position from childhood to young adulthood. Angle Orthodontist. 2009;79:484–90. doi: 10.2319/062308-328.1.

32. Mislik B, Hänggi MP, Signorelli L, Peltomäki TA, Patcas R. Pharyngeal airway dimensions: A cephalometric, growth-study-based analysis of physiological variations in children aged 6-17. European Journal of Orthodontics. 2014;36:331–9. doi: 10.1093/ejo/cjt068.

33. Daraze A, Delatte M, Liistro G, Majzoub Z. Cephalometrics of Pharyngeal Airway Space in Lebanese Adults. International Journal of Dentistry. 2017;2017. doi: 10.1155/2017/3959456.

34. Rommel N, Bellon E, Hermans R, Smet M, De Meyer AM, Feenstra L, et al. Development of the Orohypopharyngeal Cavity in Normal Infants and Young Children. Cleft Palate-Craniofacial Journal. 2003;40:606–11. doi: 10.1597/1545-1569(2003)040<0606:DOTOCI>2.0.CO;2.

35. Chiang CC, Jeffres MN, Miller A, Hatcher DC. Three-dimensional airway evaluation in 387 subjects from one university orthodontic clinic using cone beam computed tomography. Angle Orthod. 2012;82(6):985–92. Epub 2012/06/07. doi: 10.2319/122811-801.1. PubMed PMID: 22668315.

36. Jiang YY, Xu X, Su HL, Liu DX. Gender-related difference in the upper airway dimensions and hyoid bone position in Chinese Han children and adolescents aged 6-18 years using cone beam computed tomography. Acta Odontologica Scandinavica. 2014;73:391–400. doi: 10.3109/00016357.2014.978366.

37. Lenza MG, De MM, Dalstra M, Melsen B, Cattaneo PM. An analysis of different approaches to the assessment of upper airway morphology: A CBCT study. Orthodontics and Craniofacial Research. 2010;13(2):96–105. Epub 2010/05/19. doi: 10.1111/j.1601-6343.2010.01482.x. PubMed PMID: 20477969.

38. Schendel SA, Jacobson R, Khalessi S. Airway growth and development: A computerized 3-dimensional analysis. Journal of Oral and Maxillofacial Surgery. 2012;70:2174–83. doi: 10.1016/j.joms.2011.10.013.

39. Anandarajah S, Dudhia R, Sandham A, Sonnesen L. Risk factors for small pharyngeal airway dimensions in preorthodontic children: A three-dimensional study. Angle Orthod. 2017;87(1):138–46. Epub 2016/11/02. doi: 10.2319/012616-71.1. PubMed PMID: 27304232.

40. Masoud AI, Alwadei FH, Alwadei AH, Lin EY, Viana MGC, Kusnoto B, et al. Developing pediatric three-dimensional upper airway normative values using fixed and interactive thresholds. Oral Radiology. 2020;36:89–99. doi: 10.1007/s11282-019-00384-3.

41. Yanagita N, Terajima M, Kanomi R, Takahashi I. Three-dimensional analysis of pharyngeal airway morphology in Japanese female adolescents. Orthodontic Waves. 2019;76(2):89–96. doi: 10.1016/j.odw.2017.01.004.

42. Alves M, Franzotti ES, Baratieri C, Nunes LKF, Nojima LI, Ruellas ACO. Evaluation of pharyngeal airway space amongst different skeletal patterns. International Journal of Oral and Maxillofacial Surgery. 2012;41:814–9. doi: 10.1016/j.ijom.2012.01.015.

43. Claudino LV, Mattos CT, Ruellas ACDO, Sant Anna EF. Pharyngeal airway characterization in adolescents related to facial skeletal pattern: A preliminary study. American Journal of Orthodontics and Dentofacial Orthopedics. 2013. doi: 10.1016/j.ajodo.2013.01.015. PubMed PMID: 23726330.

44. Li H, Lu X, Shi J, Shi H. Measurements of normal upper airway assessed by 3-dimensional computed tomography in Chinese children and adolescents. International Journal of Pediatric Otorhinolaryngology. 2011;75:1240–6. doi: 10.1016/j.ijporl.2011.06.022.

45. Kim EJ, Choi JH, Kim YS, Kim TH, Lee SH, Lee HM, et al. Upper airway changes in severe obstructive sleep apnea: Upper airway length and volumetric analyses using 3D MDCT. Acta Oto-Laryngologica. 2011;131:527–32. doi: 10.3109/00016489.2010.535561.

46. Gibelli D, Cellina M, Gibelli S, Oliva AG, Termine G, Sforza C. Three-Dimensional Assessment of Pharyngeal Volume on Computed Tomography Scans: Applications to Anesthesiology and Endoscopy. Journal of Craniofacial Surgery. 2020;00:1. doi: 10.1097/scs.0000000000006094.

47. Inamoto Y, Saitoh E, Okada S, Kagaya H, Shibata S, Baba M, et al. Anatomy of the larynx and pharynx: Effects of age, gender and height revealed by multidetector computed tomography. Journal of Oral Rehabilitation. 2015;42:670–7. doi: 10.1111/joor.12298.

48. Shigeta Y, Takumi O, Venturin J, Nguyen M, Clark GT, Enciso R. Gender and age-based differences in computed-tomography measurements of the orophaynx. Oral Surg Oral Med Oral Pathol Oral Radiol Endod. 2008;106(4).

49. Welch KC, Foster GD, Ritter CT, Wadden TA, Arens R, Maislin G, et al. A Novel Volumetric Magnetic Resonance Imaging Paradigm to Study Upper Airway Anatomy. Sleep. 2002;25:530–40. doi: 10.1093/sleep/25.5.530.

50. Leboulanger N, Louis B, Fodil R, Boelle PY, Clement A, Garabedian EN, et al. Analysis of the pharynx and the trachea by the acoustic reflection method in children: a pilot study. Respir Physiol Neurobiol. 2011;175(2):228–33. Epub 2010/11/30. doi: 10.1016/j.resp.2010.11.008. PubMed PMID: 21111847.

51. Martin SE, Mathur R, Marshall I, Douglas NJ. The effect of age, sex, obesity and posture on upper airway size. European Respiratory Journal. 1997;10:2087–90. doi: 10.1183/09031936.97.10092087.

52. Brooks LJ, Strohl KP. Size and mechanical properties of the pharynx in healthy men and women. American Review of Respiratory Disease. 1992;146:1394–7. doi: 10.1164/ajrccm/146.6.1394.

53. Brown IG, Zamel N, Hoffstein V. Pharyngeal cross-sectional area in normal men and women. Journal of Applied Physiology. 1986;61:890–5. doi: 10.1152/jappl.1986.61.3.890.

54. Gurani SF, Di Carlo G, Cattaneo PM, Thorn JJ, Pinhol EM. Effect of Head and Tongue Posture on the Pharyngeal Airway Dimensions and Morphology in Three-Dimensional Imaging: a Systematic Review. Journal of Oral and Maxillofacial Research. 2016;7:1–12. doi: 10.5037/jomr.2016.7101.

55. Di Carlo G, Gurani SF, Pinholt EM, Cattaneo PM. A new simple three-dimensional method to characterize upper airway in orthognathic surgery patient. Dentomaxillofacial Radiology. 2017;46. doi: 10.1259/dmfr.20170042.

56. Ayappa I, Rapoport DM. The upper airway in sleep: Physiology of the pharynx. Sleep Medicine Reviews. 2003;7:9–33. doi: 10.1053/smrv.2002.0238. PubMed PMID: 12586528.

57. Adewale L. Anatomy and assessment of the pediatric airway. Paediatric Anaesthesia. 2009;19:1–8. doi: 10.1111/j.1460-9592.2009.03012.x.

58. Netter FH. Head and Neck. Atlas of Human Anatomy. 7 ed. Philadelphia, PA: Elsevier Inc.; 2019. p. 11-174.

59. Arens R, Marcus CL. Pathophysiology of upper airway obstruction: A developmental perspective. Sleep. 2004;27:997–1019. doi: 10.1093/sleep/27.5.997. PubMed PMID: 15453561.

60. Logan BM, Reynolds PA, Rice S, Hutchings RT. Nose, oral cavity, pharynx, ear and larynx. In: Logan BM, Reynolds PA, Rice S, editors. McMinn’s Color Atlas of Head and Neck Anatomy. 5 ed: Elsevier Ltd; 2017.

61. Schuenke M, Schulte E, Schumacher U. Oral Cavity and Perioral Region. In: Baker EW, editor. Head and Neck Anatomy for Dental Medicine. New York, NY: Thieme; 2010. p. 216.

62. Laird AM, Yetkiner E, Kadioglu O, Currier GF. The Upper Airway. In: Kadioglu O, Currier GF, editors. Craniofacial 3D Imaging: Current Concepts in Orthodontics and Oral and Maxillofacial Surgery. Cham: Springer International Publishing; 2019. p. 97–112.

63. Moore K, Dalley A, Agur AMR. Neck. Clinically Oriented Anatomy. 7 ed: Lippincott Williams & Wilkins; 2006.

64. Standring S. Gray’s anatomy : the anatomical basis of clinical practice. Forty-first edition. ed. New York: Elsevier Limited; 2016. p. xviii, 1562 pages.

65. Gu M, McGrath CPJ, Hägg U, Wong RWK, Yang Y. Anatomy of the Upper Airway and Its Growth in Childhood. Journal of Dentistry and Oral Biology. 2016;1.

66. Kelly MP, Vorperian HK, Wang Y, Tillman KK, Werner HM, Chung MK, et al. Characterizing mandibular growth using three-dimensional imaging techniques and anatomic landmarks. Archives of Oral Biology. 2017;77:27–38. doi: 10.1016/j.archoralbio.2017.01.018.

67. Miller CA, Hwang SJ, Cotter MM, Vorperian HK. Cervical vertebral body growth and emergence of sexual dimorphism: a developmental study using computed tomography. Journal of Anatomy. 2019;234:764–77. doi: 10.1111/joa.12976.

68. Miller CA, Lee Y, Avey GD, Vorperian HK. Head position classification of medical imaging studies: an assessment and development of a protocol. Dentomaxillofacial Radiology. 2019:20190220. doi: 10.1259/dmfr.20190220.

69. AnalyzeDirect . Analyze 12.0. Overland Park, KS: AnalyzeDirect; 2018.

70. Nakano H, Mishima K, Ueda Y, Matsushita A, Suga H, Miyawaki Y, et al. A new method for determining the optimal CT threshold for extracting the upper airway. Dentomaxillofacial Radiology. 2013;42. doi: 10.1259/dmfr/26397438.

71. Vorperian HK, Wang S, Michael Schimek E, Durtschi RB, Kent RD, Gentry LR, et al. Developmental sexual dimorphism of the oral and pharyngeal portions of the vocal tract: An imaging study. Journal of Speech, Language, and Hearing Research. 2011;54:995–1010. doi: 10.1044/1092-4388(2010/10-0097).

72. Lorensen WE, Cline HE. Marching cubes: A high resolution 3D surface construction algorithm. Proceedings of the 14th annual conference on Computer graphics and interactive techniques. 37422: ACM; 1987. p. 163–9.

73. Desbrun M, Meyer M, Schröder P, Barr AH, editors. Implicit fairing of irregular meshes using diffusion and curvature flow. Proceedings of the 26th annual conference on Computer graphics and interactive techniques - SIGGRAPH ’99; 1999.

74. Lazarus F, Verroust A. Level set diagrams of polyhedral objects. Proceedings of the Symposium on Solid Modeling and Applications. 1999:130–40. doi: 10.1145/304012.304025.

75. Seo S, Chung MK, Whyms BJ, Vorperian HK, editors. Mandible shape modeling using the second eigenfunction of the Laplace-Beltrami operator. Medical Imaging 2011: Image Processing; 2011.

76. Shi Y, Lai R, Krishna S, Sicotte N, Dinov I, Toga AW. Anisotropic Laplace-Beltrami Eigenmaps: Bridging Reeb Graphs and Skeletons. Proc IEEE Comput Soc Conf Comput Vis Pattern Recognit. 2008;2008:1–7. Epub 2008/07/15. doi: 10.1109/CVPRW.2008.4563018. PubMed PMID: 21339850; PubMed Central PMCID: PMCPMC3041984.

77. de Boor C. A Practical Guide to Splines. New York, NY: Springer New York; 1978.

78. Hunyandi L. B-splines. MATLAB Central File Exchange; 2010.

79. Hudgins PA, Siegel J, Jacobs I, Abramowsky CR. The normal pediatric larynx on CT and MR. AJNR Am J Neuroradiol. 1997;18(2):239–45. Epub 1997/02/01. PubMed PMID: 9111658.

80. Scammon RE. The measurement of the body in childhood. In: Harris JA, Jackson CM, Paterson DG, Scammon RE, editors. The measurement of man. Minneapolis, MN: The University of Minnesota Press; 1930. p. 173–215.

81. Karia H, Shrivastav S, Karia AK. Three-dimensional evaluation of the airway spaces in patients with and without cleft lip and palate: A digital volume tomographic study. Am J Orthod Dentofacial Orthop. 2017;152(3):371–81. Epub 2017/09/03. doi: 10.1016/j.ajodo.2016.12.026. PubMed PMID: 28863918.

82. Cassano P, Gelardi M, Cassano M, Fiorella ML, Fiorella R. Adenoid tissue rhinopharyngeal obstruction grading based on fiberendoscopic findings: a novel approach to therapeutic management. Int J Pediatr Otorhinolaryngol. 2003;67(12):1303–9. Epub 2003/12/04. doi: 10.1016/j.ijporl.2003.07.018. PubMed PMID: 14643473.

83. Jeans WD, Fernando DCJ, Maw AR, Leighton BC. A longitudinal study of the growth of the nasopharynx and its contents in normal children. The British Journal of Radiology. 1981;54(638):117–21. doi: 10.1259/0007-1285-54-638-117. PubMed PMID: 7459548.

84. de Souza Vilella B, de Vasconcelos Vilella O, Koch HA. Growth of the nasopharynx and adenoidal development in Brazilian subjects. Brazilian Oral Research. 2006;20:70–5. doi: 10.1590/s1806-83242006000100013.

85. Linder-Aronson S, Stockholm BCL, London S, editors. A longitudinal study of the development of the posterior nasopharyngeal wall between 3 and 16 years of age. European Journal of Orthodontics; 1983.

86. Luscan R, Leboulanger N, Fayoux P, Kerner G, Belhous K, Couloigner V, et al. Developmental changes of upper airway dimensions in children. Pediatric Anesthesia. 2020:pan.13832. doi: 10.1111/pan.13832. PubMed PMID: 31995659.

87. Liu S, Qi W, Zhang X, Dong Y. The development of the cricoid cartilage and its implications for the use of endotracheal tubes in the pediatric population. Pediatric Anesthesia. 2020;30(1):63–8. doi: https://doi.org/10.1111/pan.13772.

88. Perry JL, Kollara L, Kuehn DP, Sutton BP, Fang X. Examining age, sex, and race characteristics of velopharyngeal structures in 4-to 9-year-old children using magnetic resonance imaging. Cleft Palate-Craniofacial Journal. 2018;55:21–34. doi: 10.1177/1055665617718549.

89. Dang J, Honda K. Acoustic Characteristics of the Piriform Fossa in Models and Humans. Journal of the Acoustical Society of America. 1997;101(1):456–65. doi: 10.1121/1.417990.

90. Story BH. Physiologically-based speech simulation using an enhanced wave-reflection model of the vocal tract [Ph.D.]. Ann Arbor: The University of Iowa; 1995.

91. Story BH, Titze IR, Hoffman EA. Vocal tract area functions for an adult female speaker based on volumetric imaging. The Journal of the Acoustical Society of America. 1998;104(1):471–87.

92. Fant G. Acoustic Theory of Speech Production. Acoustic Theory of Speech Production. 1971. doi: 10.1515/9783110873429.

93. Baer T, Gore JC, Gracco LC, Nye PW. Analysis of Vocal Tract Shape and Dimensions using Magnetic Resonance Imaging: Vowels. Journal of the Acoustical Society of America. 1991;90(2 Pt 1):799–828.

94. Fujita S, Honda K. An experimental study of acoustic characteristics of hypopharyngeal cavities using vocal tract solid models. Acoustical Science and Technology. 2005;26(4):353–7. doi: 10.1250/ast.26.353.

95. Story BH, Bunton K. A model of speech production based on the acoustic relativity of the vocal tract. J Acoust Soc Am. 2019;146(4):2522. Epub 2019/11/02. doi: 10.1121/1.5127756. PubMed PMID: 31671993.

96. Buhr RD. The Emergence of Vowels in an Infant. Journal of Speech, Language, and Hearing Research. 1980;23(1):73–94.

97. Watters KF. Tracheostomy in Infants and Children. Respiratory Care. 2017;62:799–825.

98. Thach BT. The Role of the Upper Airway in SIDS and Sudden Unexpected Infant Deaths and the Importance of External Airway-Protective Behaviors. In: Duncan JR, Byard RW, editors. SIDS Sudden Infant and Early Childhood Death: The Past, the Present and the Future. Adelaide (AU)2018.

99. Cornwell AC. Sex differences in the maturation of sleep/wake patterns in high risk for SIDS infants. Neuropediatrics. 1993;24(1):8–14. Epub 1993/02/01. doi: 10.1055/s-2008-1071505. PubMed PMID: 8474614.

100. Khadivi E, Zaringhalam MA, Khazaeni K, Bakhshaee M. Distance between anterior commissure and the first tracheal ring: An important new clinical laryngotracheal measurement. Iranian Journal of Otorhinolaryngology. 2015;27(80):193–7. doi: 10.22038/ijorl.2015.4252.

101. Sirisopana M, Saint-Martin C, Wang NN, Manoukian J, Nguyen LHP, Brown KA. Novel measurements of the length of the subglottic airway in infants and young children. Anesthesia and Analgesia. 2013;117(2):462–70. doi: 10.1213/ANE.0b013e3182991d42.

102. Griscom NT, Wohl MEB. Dimensions of the growing trachea related to age and gender. American Journal of Roentgenology. 1986;146(2):233–7. doi: 10.2214/ajr.146.2.233.

103. Perry JL, Kuehn DP, Sutton BP, Gamage JK, Fang X. Anthropometric analysis of the velopharynx and related craniometric dimensions in three adult populations using MRI. Cleft Palate-Craniofacial Journal. 2016;53:e1–e13. doi: 10.1597/14-015.

104. Bloom K, Moore-Schoenmakers K, Masataka N. Nasality of infant vocalizations determines gender bias in adult favorability ratings. Journal of Nonverbal Behavior. 1999;23(3):219–36. doi: Doi 10.1023/A:1021317310745. PubMed PMID: WOS:000083422500002.

105. Yang S, Mu L. An investigation of the third formant of /a/ in prepubertal children. Journal of Voice. 1989;3(4):321–3. doi: https://doi.org/10.1016/S0892-1997(89)80054-2.

106. Perry TL, Ohde RN, Ashmead DH. The acoustic bases for gender identification from children’s voices. The Journal of the Acoustical Society of America. 2001;109(6):2988–98.

107. Vorperian HK, Kent RD, Lee Y, Bolt DM. Corner vowels in males and females ages 4 to 20 years: Fundamental and F1-F4 formant frequencies. J Acoust Soc Am. 2019;146(5):3255. Epub 2019/12/05. doi: 10.1121/1.5131271. PubMed PMID: 31795713; PubMed Central PMCID: PMCPMC6850954.

108. Buschang PH, Hinton RJ. A Gradient of Potential for Modifying Craniofacial Growth. Seminars in Orthodontics. 2005;11(4):219–26. doi: 10.1053/j.sodo.2005.07.006.

109. Wang Y, Chung MK, Vorperian HK. Composite growth model applied to human oral and pharyngeal structures and identifying the contribution of growth types. Statistical Methods in Medical Research. 2016;25:1975–90. doi: 10.1177/0962280213508849.

110. Durtschi RB, Chung D, Gentry LR, Chung MK, Vorperian HK. Developmental craniofacial anthropometry: Assessment of race effects. Clinical Anatomy. 2009;22(7):800–8. Epub 2009/09/16. doi: 10.1002/ca.20852. PubMed PMID: PMID: 19753647

